# The PELOTA-HBS1 Complex Orchestrates mRNA Translation Surveillance and PDK1-mediated Plant Growth and Development

**DOI:** 10.1101/2021.01.11.426208

**Authors:** Wei Kong, Shutang Tan, Qing Zhao, De-Li Lin, Zhi-Hong Xu, Jiří Friml, Hong-Wei Xue

**Author notes:** **Lead contact / Corresponding author:** Shanghai Jiao Tong University, 800 Dongchuan Road, Shanghai 200240, China. **Contributed equally.** (Wei Kong, Shutang Tan).

## Abstract

The quality control system for messenger RNA is fundamental for cellular activities in eukaryotes. To elucidate the molecular mechanism of 3’-Phosphoinositide-Dependent Protein Kinase1 (PDK1), an essential regulator throughout growth and development of eukaryotes, a forward genetic approach was employed to screen for suppressors of the loss-of-function T-DNA insertional *pdk1*.*1 pdk1*.*2* double mutant in *Arabidopsis*. Notably, the severe growth attenuation of *pdk1*.*1 pdk1*.*2* is rescued by *sop21* (<u>s</u>uppressor <u>o</u>f *<u>p</u>dk1*.*1 pdk1*.*2*) that harbours a loss-of-function mutation in *PELOTA1* (*PEL1*). PEL1 is a homologue of mammalian PELOTA and yeast DOM34, which form a heterodimeric complex with the GTPase HBS1, responsible for ribosome rescue to assure the quality and fidelity of mRNA molecules. Genetic analysis further reveals that the dysfunction of PEL1-HBS complex fails to degrade the T-DNA-disrupted, truncated but functional PDK1 transcripts, thus rescuing *pdk1*.*1 pdk1*.*2*. Our studies demonstrate the functionality and identify the essential functions of a homologous PELOTA-HBS1 complex in higher plant, and provide novel insights into the mRNA quality control mechanism.

## Introduction

Living organisms need to monitor both the quantity and the quality of biomolecules, such as nucleic acids and proteins, to accomplish various life activities. Protein quality control is ensured by multi-level regulations both translationally and post-translationally, of which, the messenger RNA (mRNA) quality is essential for the biosynthesis of correct corresponding proteins. The quality and fidelity of mRNAs are monitored by cells autonomously, and aberrant mRNAs need to be recognised by intrinsic molecular machineries, to release stalled ribosomes and get degraded themselves (*1*). According to current understandings, there are at least three different mechanisms for mRNA degradation: nonsense-mediated decay (NMD), no-stop decay (NSD), and no-go decay (NGD). NMD and NSD target mRNAs with premature stop codon (terminated too soon) and lacking stop codon (failing to terminate) respectively. In mammalian cells and yeasts, the PELOTA-HBS1 (Dom34p-HBS1p) complex plays an essential role in regulating NSD (*2*). However, little was known about the mRNA quality control in plants so far.

3’-Phosphoinositide-Dependent Protein Kinase1 (PDK1) is conserved in eukaryotes and plays important roles in regulating growth and development in various organisms. As a key member of the cAMP-dependent protein kinase A / protein kinase G / protein kinase C (AGC) kinase family (*3*), PDK1 is important for the activation of many AGC kinases and other substrates / regulators. Studies have revealed that PDK1 plays crucial roles in the signalling pathways activated by growth factors and hormones, sustains and regulates the balance between cell growth, division and apoptosis in mammals (*3*-*5*), thus being critical for normal development. However, loss-of-function mutant *pdk1* in various species such as yeasts (*6, 7*), *Drosophila* (*8*) and mice (*9*), is lethal, which makes it challenging to study the downstream regulations, and functional mechanism of PDK1 is still not completely understood yet.

Differently from those of mammals, loss-of-function or knock-down *pdk1* mutant plants are viable, despite exhibiting severe developmental defects, including rice (*10*), *Arabidopsis* (*11*–*13*) and moss (*14*). Therefore, it provides a plausible approach to further identify genetic interactors of PDK1. There are two PDK1 paralogues in *Arabidopsis*, PDK1.1 and PDK1.2, which have redundant functions (*12, 15*). PDK1 binds to phospholipids, which regulate its activity as well as its subcellular localization (*12, 16, 17*). As a master regulator of the AGC family, PDK1 was proposed to participate in various growth and developmental processes though phosphorylating distinct kinase substrates (*12, 18*). For example, PDK1.1 regulates root hair development through phosphorylating OXIDATIVE SIGNAL INDUCIBLE1 (OXI1)/AGC2-1 kinase (*19, 20*). PINOID (PID), an essential regulator of PIN FORMED (PIN) auxin efflux carriers, was phosphorylated by PDK1 and thus being activated *in vitro* (*21, 22*). Recently, characterization of the fully knock-out *pdk1*.*1 pdk1*.*2* double mutant, uncovers the important role of plant PDK1. Both PDK1.1 and PDK1.2 are expressed in vascular tissues, and show a predominant localization at the basal side of cell plasma membrane (PM) as well as at cytoplasm in root stele. Notably, the *pdk1*.*1 pdk1*.*2* double mutant has pleotropic defects throughout growth and development, revealing an essential function of PDK1 in divergent life activities (*12, 13*). Importantly, the basal localization of PDK1 dominants the role of these AGC kinases with the same subcellular distribution, including D6 Protein Kinase (D6PK) / D6 Protein Kinase Likes (D6PKLs) (*23*) and PROTEIN KINASE ASSOCIATED WITH BRX (PAX) (*24*), and thus participate in the regulation of polar auxin transport (*12, 13, 15*).

To further identify regulators involved in the PDK1 pathway, a forward genetic approach was employed. Using an EMS population of *Arabidopsis pdk1*.*1 pdk1*.*2* double mutant that displays severe growth defects, a suppressor screening was performed. In this study, characterization of the identified mutant, *sop21* (suppressor of *pdk1*.*1 pdk1*.*2*), reveals that deficiency of translational mRNA surveillance PELOTA-HBS1 complex rescues the defective phenotype of *pdk1*.*1 pdk1*.*2*. Our studies demonstrate the functionality of a homologous PELOTA-HBS1 complex in higher plants and provide informative clues on the control of mRNA surveillance and thus protein homeostasis.

## Results

### Deficiency of *PEL1* suppresses the defective growth of *Arabidopsis pdk1.1 pdk1.2*

PDK1 is well known as a master regulator of AGC family in eukaryotic kingdom. Two *PDK1* paralogous genes in *Arabidopsis thaliana, PDK1.1* (AT5G04510) and *PDK1.2* (AT3G10540) (*11, 12, 14*), exhibit overlapping and widespread expression pattern in various tissues (Supplementary Fig. 1). The *pdk1* single loss-of-function mutants, *pdk1.1-2* and *pdk1.2-4* (hereafter as *pdk1.1* and *pdk1.2* respectively), displayed no obvious growth phenotype (*12*), whereas, the double mutant *pdk1.1 pdk1.2* exhibits a range of developmental defects (Fig. 1, and Supplementary Fig. 2), including suppressed growth (smaller leaves and shorter siliques), reduced axillary shoots, suppressed primary root elongation and lateral root initiation, particularly significantly decreased fertility, and abnormal floral development (*12*). This is consistent with the crucial roles of PDK1 in other organisms and confirms the essential role of *PDK1* in regulating plant growth and development. Several aspects of those developmental defects can be explained by known AGC kinases, such as D6PK (*12*), PAX (*13*), and AGC1.5/7 (*13, 25*). However, whether there are additional components in the PDK1 pathway, other than AGC kinases, remains unclear.

**Fig 1.**
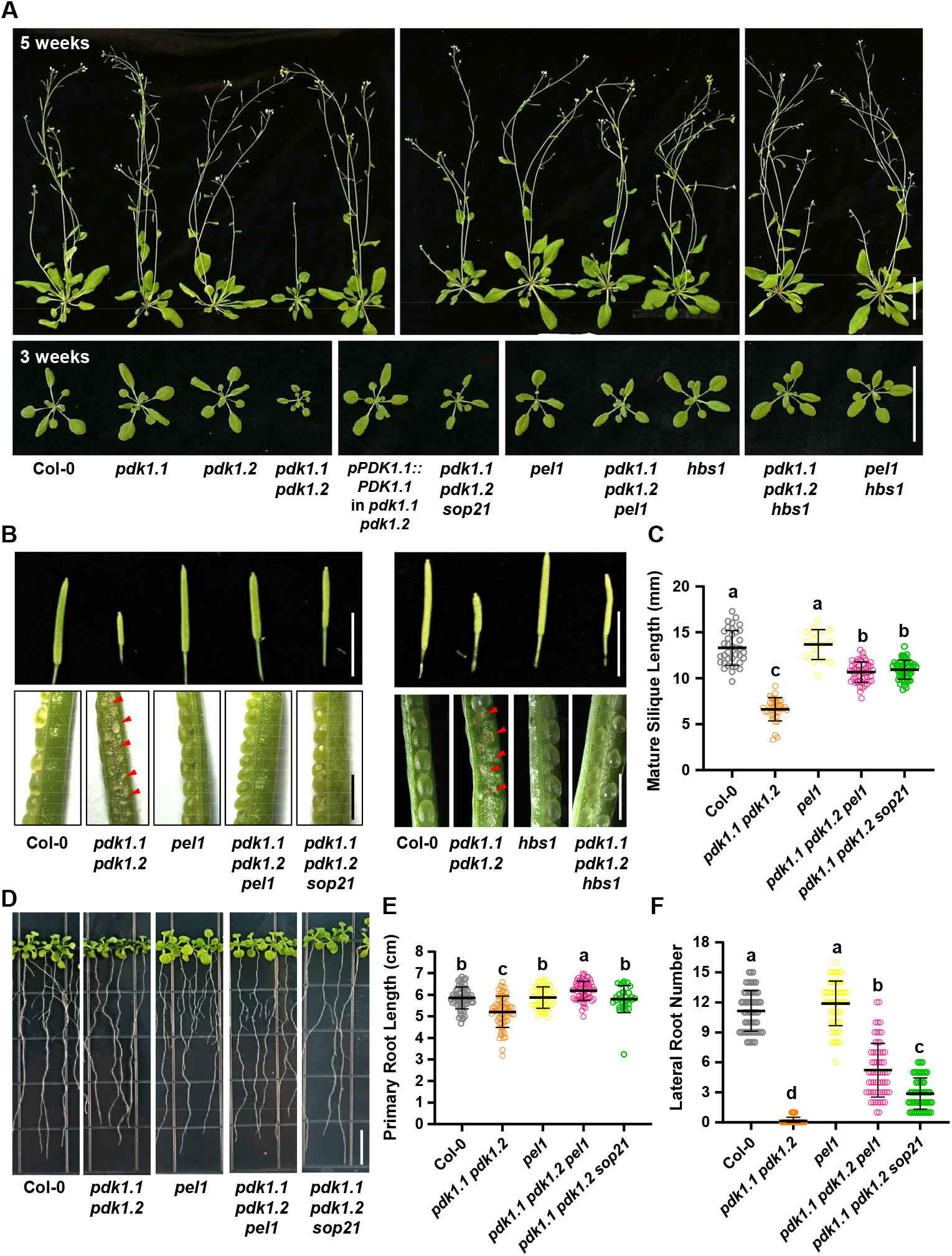
The *sop21* mutation or deficiency of *PEL1 or HBS1* restores the growth defects of *Arabidopsis pdk1*.*1 pdk1*.*2* mutants. A. The *sop21* mutation or loss of function of *PEL1* or *HBS1* suppresses the growth defect of *pdk1.1 pdk1.2*, including the reduced growth and delayed bolting. 3- or 5-week-old Col-0, *pdk1.1 pdk1.2, pdk1.1 pdk1.2 pel1, pdk1.1 pdk1.2 sop21* (*pdk1.1 pdk1.2 pel1*^*W27\**^), *pdk1.1 pdk1.2 hbs1* and various mutant plants are shown. Scale bar, 5 cm. b. Short siliques and low setting rate, Scale bars, 2 cm (upper) or 1 mm (lower), defective/abnormal seeds of *pdk1.1 pdk1.2* are highlighted. B. Quantification of the silique length. n = 37, 36, 17, 54, and 51, respectively. Different letters represent significant difference, *P <* 0.05, by one-way analysis of variance (ANOVA) with a Tukey multiple comparison test. C. Defective root elongation and lateral root formation, 2-week-old seedlings, Scale bar, 1 cm). Representative images are shown. D. Length of primary root and number of emerged lateral roots of 2-week-old seedlings were calculated. Data are presented as means ± SD (n > 30). n = 56, 65, 52, 62, and 30, respectively. Different letters represent significant difference, *P <* 0.05, by one-way analysis of variance (ANOVA) with a Tukey multiple comparison test. E. Number of emerged lateral roots of two-week-old seedlings were counted. and statistically analyzed by student’s *t-*test (**, p < 0.01). Data are presented as means ± SD (n > 30). n = 61, 65, 63, 63, and 52, respectively. Different letters represent significant difference, *P <* 0.01, by one-way ANOVA with a Tukey multiple comparison test.

To elucidate the underlying mechanism of PDK1 function, a forward genetic screen was performed. Seeds of *pdk1.1 pdk1.2* were used to generate a mutant population by Ethyl methanesulfonate (EMS) mutagenesis and suppressors of *pdk1.1 pdk1.2* (*sop, suppressor of pdk1.1 pdk1.2*) were screened based on the rescued growth (*26*). More than 10 suppressors were obtained from a M_2_ population of approximately 80,000 plants, and a recessive mutant, *sop21*, that showed an obviously rescued growth of *pdk1.1 pdk1.2* (Fig. 1 and Supplementary Fig. 2), was characterized first. Compared to a significantly suppressed growth of *pdk1.1 pdk1.2* adult plants, the rosette size of *pdk1.1 pdk1.2 sop21* is comparable to that of wild type (WT, Fig. 1A and Supplementary Fig. 2D). Similar degree of rescue was also observed for the fertility, silique length, inflorescence morphology and floral development (Fig. 1B, C, and Supplementary Fig. 2E, F). Notably, the lateral root numbers was only partially rescued to approximately 50% of WT, though with a complete rescue of primary root growth (Fig. 1D, E, F).

To identify the causative mutation in *sop21*, 102 progenies (referred as BC_1_F_2_) showing rescue phenotypes were selected from a segregating pool of F_2_ individuals [102 rescued phenotype: 296 *pdk1.1 pdk1.2* phenotype, for 1:3 ratio (χ^2^=0.084, P>0.75; Chi-square test), which indicated a single recessive causal mutation] and used for DNA extraction and subsequent deep sequencing (*27*). Systemic analysis revealed that *sop21* carried a mutation in *PEL1* (AT4G27650) (*28*), which led to an early stop at amino acid residue 27 (tryptophan to terminator, W27*), resulting in the translationally premature termination (Supplementary Fig. 2A). *PEL1* gene is widespread expressed (Supplementary Fig. 3) and cross of a null T-DNA insertional allele *pel1* (SALK_124403, also named *lesion mimic leaf1-1, lml1-1* (*29*), Supplementary Fig. 2B, C) with *pdk1.1 pdk1.2* also suppressed the growth defects of *pdk1.1 pdk1.2* at various aspects (Fig. 1), verifying that suppression of *pdk1.1 pdk1.2* phenotype in *sop21* was a result of *PEL1* deficiency. Though *pel1* mutant was previously shown to exhibit a delayed growth rate (*29*), both *sop21* and *pel1* mutants grew normally in our hands, perhaps due to different growth conditions. In addition, expression of *PEL1-FLAG* driven by a *CaMV35S* promoter in *sop21* restored the *pdk1.1 pdk1.2* phenotype (Supplementary Fig. 4A, B), confirming that *PEL1* deficiency suppressed the growth defects of *pdk1.1 pdk1.2*. Overexpression of *PEL1-FLAG* driven by *CaMV35S* in WT background did not exhibit any obvious phenotype (Supplementary Fig. 4C, D).

Next, we constructed *35S::PEL1-GFP* and *35S::mCherry-PEL1* transgenic plants to study the subcellular localization of PEL1. Presence of PEL1-GFP and mCherry-PEL1 in cytoplasm and nucleus was observed (Fig. 2A). The subcellular localization of PEL1 is partially overlapping with that of PDK1.1, which also showed residence at cytoplasm, except for its PM localization (Fig. 2A). Further analysis using tobacco leaves revealed that both mCherry-PEL1 (Fig. 2B) and mCherry-PDK1.1 (Fig. 2C) did not distribute equally in cytoplasm, and indeed they exhibited a similar feature that both proteins localized to certain compartments associated with endoplasmic reticulum (ER).

**Fig 2.**
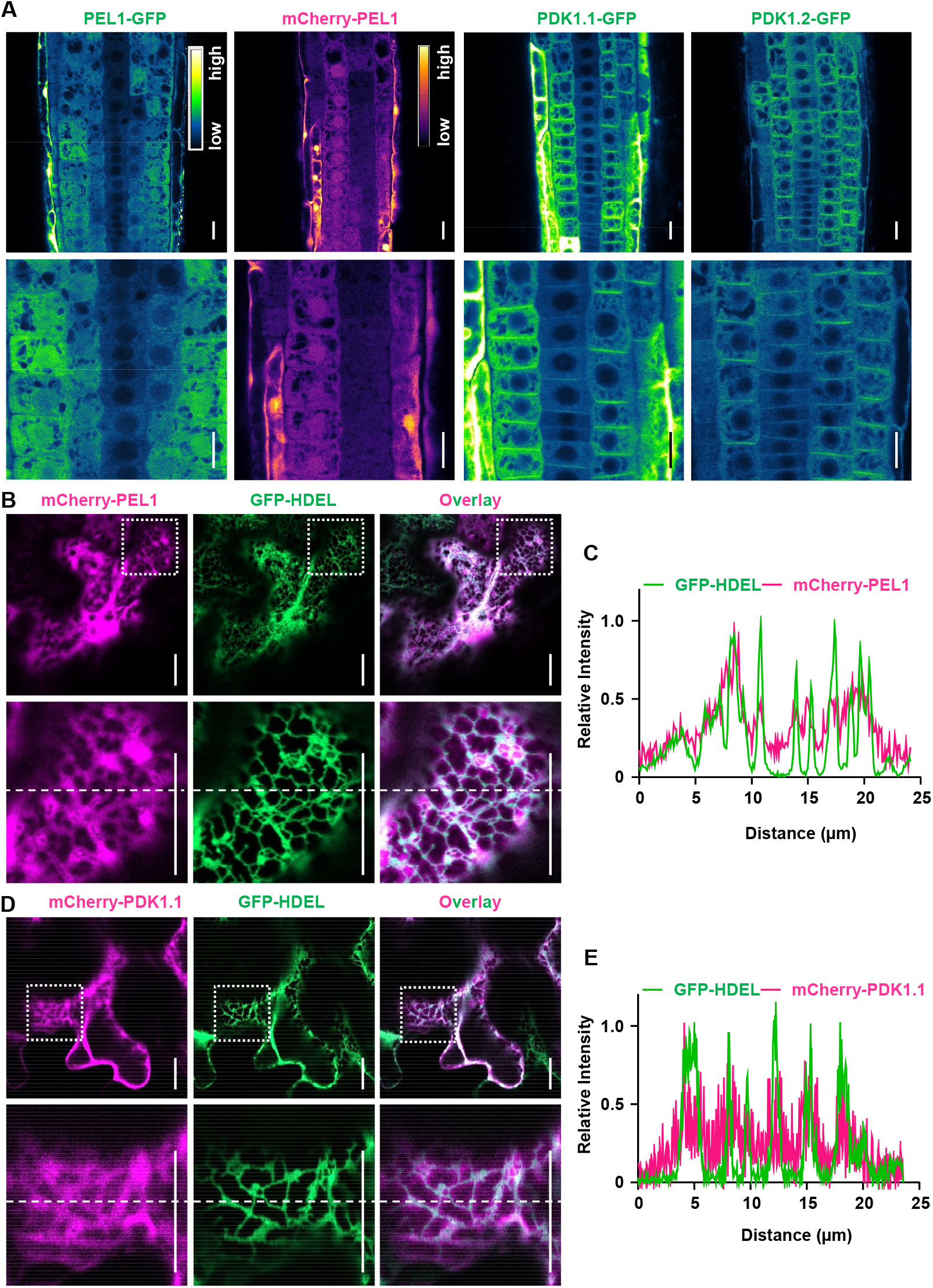
Subcellular localizations of PEL1 and PDK1.1. A. Stable transgenic lines revealed that PEL1 localized to the cytoplasm and nucleus, and that PDK1.1-GFP and PDK1.2-GFP resided at PM and cytoplasm. Five-day-old *35S::PEL1-GFP, 35S::mCherry-PEL1, 35S::PDK1.1-GFP* and *35S::PDK1.2-GFP* seedlings were observed by CLSM. The “Green Fire Blue” LUT was used for GFP, and “mpl-inferno” LUT was used for mCherry, visualizations respectively, based on fluorescence intensity by Fiji. Scale bars, 20 µm. B-E. Fluorescence observations showed that both PEL1 (b, c) and PDK1.1 (d, e) localized to certain cytoplasm compartments associated with the endoplasmic reticulum (ER). Fusion proteins PEL1-GFP (b, c) and PDK1.1-GFP (d, e) were transiently expressed with ER-specific GFP-HDEL proteins in tobacco leaves. Samples were observed 48 hours after infiltration. Scale bars, 20 μm. Lower panels are enlarged view of the squared region of the upper panels. The position for quantification (right panels) was indicated with dashed lines across the images.

### Presence of the mRNA surveillance complex PELOTA-HBS1 in *Arabidopsis*

Translation of aberrant mRNAs leads to stalling of translational machinery, and arrested ribosomes are specifically recognized by the PELOTA-HBS1 complex to initiate their recycling (*1, 2, 30, 31*). Mammalian and yeast PELOTA/DOM34 interacts with the HBS1 GTPase, a translation elongation factor EF1A/initiation factor IF2γ family protein, to form a heterodimer and bind to stalled ribosomes, ultimately leading to ribosome rescue. Besides, HBS1 (also called SKI7) is also involved in post-transcriptional gene silencing (*28, 32*). Homologous analysis in *Arabidopsis* genome by using yeast and human HBS1 identified two candidate HBS1 homologues, AT5G10630 and AT1G18070 (Table S1. Sequence alignment analysis showed that only AT5G10630 had the conserved HBS1_C domain, indicating that AT5G10630 (designated as AtHBS1, HBS1) is the HBS1 homologue in *Arabidopsis* (Supplementary Fig. 5). Considering that HBS1 functions in the same pathway as PELOTA(*28*), which is then chosen for further investigations. Indeed, analysis through yeast two-hybrid (Fig. 3A), bimolecular fluorescence complementation (BiFC, Fig. 3B) and GST pull-down assays (Fig. 3C) revealed the PEL1-HBS1 interactions both *in vivo* and *in vitro*, confirming the presence of a homologous PELOTA-HBS1 complex in higher plants. In addition, a recent study characterizing the roles of PELOTA and HBS1 in nonstop mRNA decay (*28*) further supports the presence of a functional PELOTA-HBS1 complex in *Arabidopsis*.

**Fig 3.**
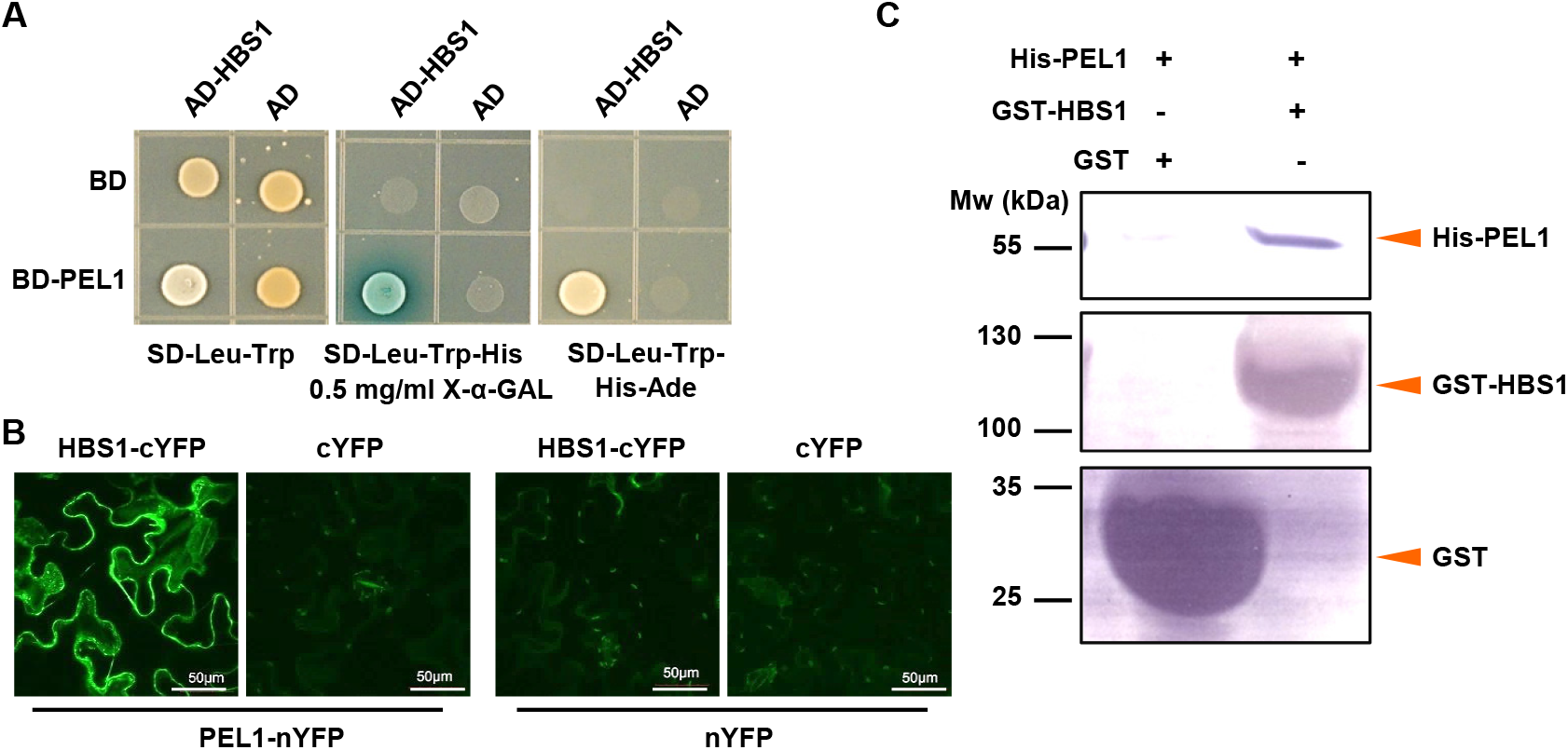
PEL1 forms a complex with HBS1. A-B. Yeast two-hybrid (A) and bimolecular fluorescence complementation (BiFC, B) analysis reveals the interactions of PEL1 with HBS1. PEL1 and HBS1 was fused to GAL4 DNA-binding domain (BD) or activation domain (AD) respectively. Protein interaction was examined on synthetic dropout (-Leu/-Trp/-His) medium supplemented with 0.5 mg/ml X-α-Gal or synthetic dropout (-Leu/-Trp/-His-Ade) medium. For BiFC analysis, PEL1-nYFP or HBS1-cYFP fusion proteins were transiently expressed in *N. benthamiana* leaves through infiltration and observed. Scale bars, 50 µm. C. GST pull-down analysis reveals the interactions of PEL1 with HBS1. GST and GST-HBS1 fusion protein were used as baits, and 6XHis-PEL1 fusion protein was used as prey. Pulled-down fractions were analyzed by Western blot using anti-His and anti-GST antibodies.

Similarly as *PEL1* and *PDK1s, HBS1* gene was ubiquitously expressed (Supplementary Fig. 6A) and a knock-down mutant of *HBS1, hbs1*, (Supplementary Fig. 6B-D) also suppressed the *pdk1.1 pdk1.2* phenotypes (Fig. 1A, B), indicating that deficiency of the translational mRNA surveillance complex PELOTA-HBS1 led to the suppression of *pdk1.1 pdk1.2* phenotypes. These observations suggest that the PELOTA-HBS1 complex might function via a common mechanism.

### PELOTA-HBS complex regulates the proper expression of truncated PDK1 transcripts

The *pdk1.1-2* and *pdk1.2-4* alleles we used contained T-DNA insertions close to their 3’-ends. It has been shown by Xiao and Offringa that such alleles are likely to lead to the production of a functional truncated protein. Therefore, in the same publication this was proposed to explain the lack of strong phenotypes for double mutant combinations *pdk1.1-2 pdk1.2-2* and *pdk1.1-2 pdk1.2-3* in previous studies *(11, 33)*. It is unlikely that this is the case, as both recent studies by Tan et al. (*12*) and Xiao and Offringa (*13*) reported highly similar phenotypes for different allele combinations. Further comparisons confirmed that the *pdk1.1 pdk1.2* combination used in this study (*pdk1.1-2 pdk1.2-4*) as well as double mutants generated with *pdk1.1-2* and other *pdk1.2* T-DNA alleles recapitulated the phenotype of double mutant combinations with the CRISPR alleles *pdk1.1-13* or *pdk1.1-14* reported by Tan et al. (*12*) and Xiao and Offringa (*13*) (Supplementary Fig. 7A,B). We speculate that previous failures to observe strong growth and development defects in *pdk1.1-2 pdk1.2-2* or *pdk1.1-2 pdk1.2-3* double mutant lines (*11, 33*) are most likely due to a failure to achieve true double homozygous plants. Interestingly, we also noted that our *pdk1.1 pdk1.2* plants are larger than the other five double mutant combinations, confirming that still a low level of functional PDK1 is produced in these plants.

Interestingly, it was also noted that *pdk1.1 pdk1.2* plants are larger than other double mutant combinations, suggesting the remaining function of PDKs. RT-qPCR analysis revealed that there was no detectable expression for both *PDK1.1* and *PDK1.2* in *pdk1.1 pdk1.2* backgrounds, with the primers across T-DNA insertions (Fig. 2D, E), ruling out the possibility of existing full-length *PDK1.1* or *PDK1.2* transcripts. Nonetheless, increased truncated transcript levels of *PDK1.1* and *PDK1.2* were detected in *pdk1.1 pdk1.2 sop21, pdk1.1 pdk1.2 pel1* or *pdk1.1 pdk1.2 hbs1* (to approximately 50% and 90% of Col-0 respectively), compared to that in *pdk1.1 pdk1.2* (10% and 25% of Col-0 for *PDK1.1* and *PDK1.2* respectively), with primers amplifying the fragments before T-DNA insertions (Fig. 2F, G). This was further confirmed by semi-quantitative PCR (Supplementary Fig. 8A). The PELOTA-HBS1 complex is responsible for the release of arrested ribosomes during the translation of aberrant mRNAs, so called “mRNA surveillance” (*28*). Given that T-DNA insertions at 3’-end might lead to aberrant *PDK1.1* and *PDK1.2* transcripts fused with certain T-DNA fragments but without a proper stop codon, it is not surprising to detect only 10% and 25% of *PDK1.1* and *PDK1.2* 5’-fragments in *pdk1.1 pdk1.2*. Consistently, there was an increase of their expression in *pdk1.1 pdk1.2 sop21, pdk1.1 pdk1.2 pel1* or *pdk1.1 pdk1.2 hbs1* plants. The above observations led us to test whether the increased levels of truncated PDK1.1 or PDK1.2 could explain for the rescue of *pdk1.1 pdk1.2*.

By transforming a mCherry-fused PDK1.1N (1-480 aa) (*12*) driven by *pPDK1.1* promoter into *pdk1.1 pdk1.2*, a partial rescue was observed (Fig. 2H). In addition, overexpression of *Venus-PDK1.1N* or *Venus-PDK1.2N* driven by a *CaMV35S* promoter completely rescued the phenotype of *pdk1.1 pdk1.2* (Supplementary Fig. 8B). Therefore, we conclude that the *pel1* and *hbs1* mutations might rescue the phenotype of *pdk1.1 pdk1.2* via disrupting the function of PELOTA-HBS1 mRNA surveillance complex and thus upregulating the N-terminal truncated proteins of PDK1.1 and PDK1.2, which preserves a functional kinase activity (*13*).

### PDK1 regulates development and stress responses through coordinating multiple metabolic pathways

The PELOTA-HBS1 complex regulates the mRNA quality control by rescuing stalled ribosomes during protein biosynthesis (*28*). PDK1 is also an essential regulator of protein translation, via modulating the activity of ribosome RPS6 proteins through S6K AGC kinase (*34*). We therefore performed a proteomic analysis to study their functions at the whole proteome level. First, being consistent, subcellular localization analysis by transiently expressing YFP- or GFP-fused proteins in *Arabidopsis* leaf protoplasts and tobacco leaves clearly showed that PDK1, PEL1 and HBS1 proteins located at plasma membrane (PM), cytoplasm and certain ER associated compartments at cytoplasm (Fig. 2B; Supplementary Fig. 9). In addition, PEL1-YFP exhibits nuclear distribution as well, which was undetectable for PDK1s and HBS1 (Fig. 2A; Supplementary Fig. 9A, B). We speculate that these proteins might present differential functions beyond the potential common pathways.

A tandem mass tag (TMT)-based comparative proteomics analysis was then performed using shoots and roots of two-week-old seedlings, and 6995 and 8137 proteins were quantified in shoots and roots respectively. We studied shoots and roots separately, because they might have totally different proteomes. Of the identified proteins, 54 and 49 proteins in shoots or roots respectively (one protein in both shoots and roots, Fig. 5A) were significantly changed in *pdk1.1 pdk1.2*, while rescued (no significant difference from WT) in *pdk1.1 pdk1.2 pel1*. These changed proteins were designated as RCE (<u>r</u>estored <u>c</u>ommonly <u>e</u>xpressed) proteins and were speculated being responsible for the defective growth of *pdk1.1 pdk1.2*. Most RCE proteins showed increased levels (84 of 102 proteins) in *pdk1.1 pdk1.2*, suggesting that PDK1 deficiency led to the enhanced recycling of ribosomes and hence the increased abundance of RCE proteins, further confirming that PDK1-mediated regulation of PELOTA-HBS1 complex is crucial to maintain the normal recycling of ribosomes and protein synthesis.

**Fig 4.**
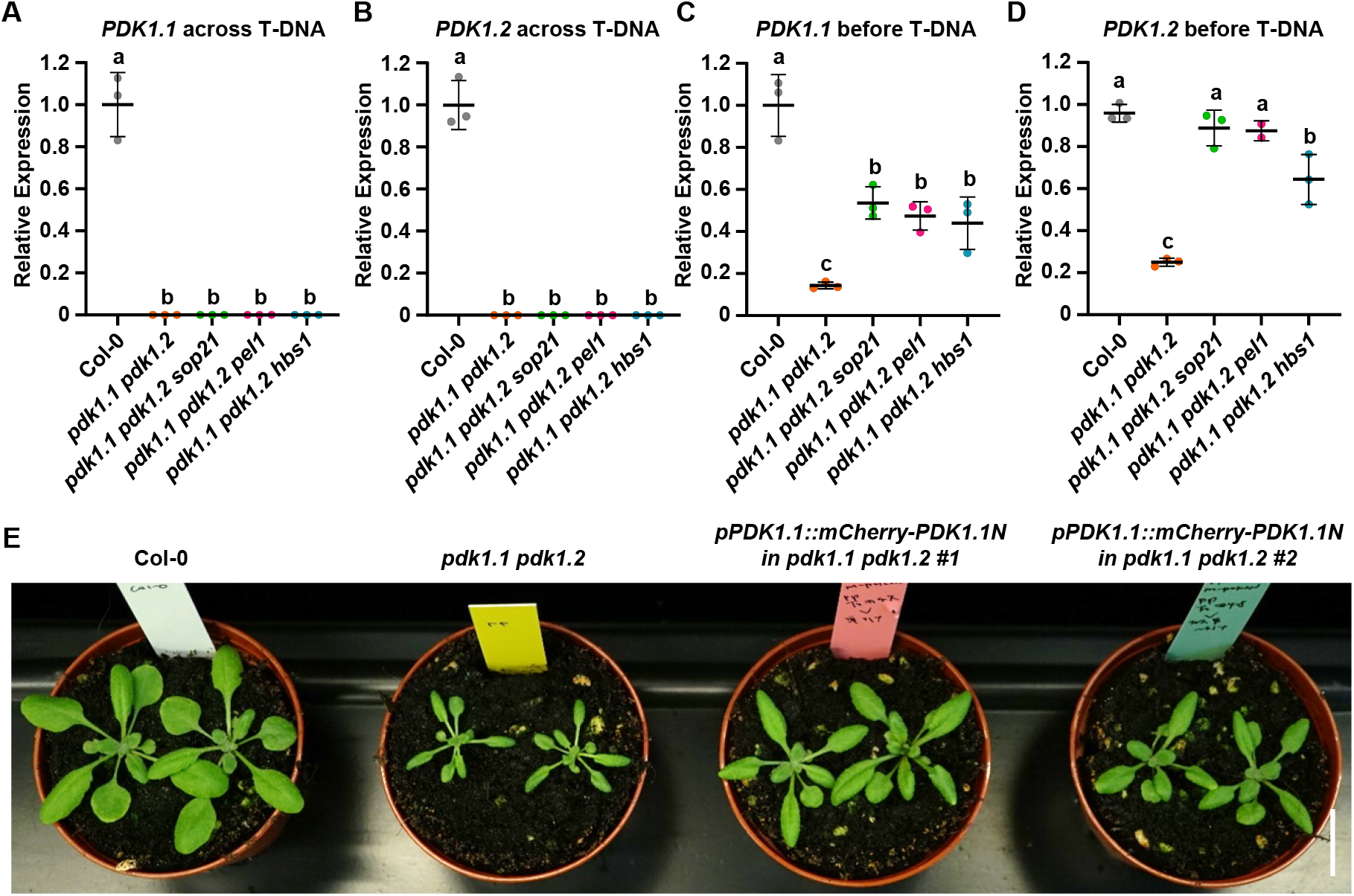
Increased expression of truncated *PDK1* transcripts in the *pdk1*.*1 pdk1*.*2 pel1* or *pdk1*.*1 pdk1*.*2 hbs1* background accounts for the rescued phenotypes. A-B. RT-qPCR analysis with primers across T-DNA insertions revealed that the integrity of *PDK1.1* and *PDK1.2* full-length CDS was disrupted by the T-DNA insertions in *pdk1.1 pdk1.2, pdk1.1 pdk1.2 sop21, pdk1.1 pdk1.2 pel1* and *pdk1.1 pdk1.2 hbs1*, respectively. *ACTIN7* gene was amplified and used as an internal control. Experiments were biologically repeated 3 times and data are presented as means ± SD. n = 3. Different letters represent significant difference, *P <* 0.05, by one-way ANOVA with a Tukey multiple comparison test. C-D. RT-qPCR analysis with primers in front of T-DNA insertions revealed that N-terminal fragments of *PDK1.1* and *PDK1.2* transcripts (PDK1.1N and PDK1.2N) exhibited increased levels in *pdk1.1 pdk1.2 sop21, pdk1.1 pdk1.2 pel1* and *pdk1.1 pdk1.2 hbs1*, respectively, compared to that in *pdk1.1 pdk1.2. ACTIN7* gene was used as an internal control. Experiments were biologically repeated 3 times and data are presented as means ± SD. n = 3. Different letters represent significant difference, *P <* 0.05, by one-way ANOVA with a Tukey multiple comparison test. E. Native promoter-driven expression of PDK1 N-terminal fragment partially rescued the growth defects of *pdk1.1 pdk1.2*. A representative photo of 20-day-old Col-0, *pdk1.1 pdk1.2*, and *pPDK1.1::mCherry-PDK1.1N* (in *pdk1.1 pdk1.2*) plants grown in soil are shown. Scale bar, 2 cm.

**Fig 5.**
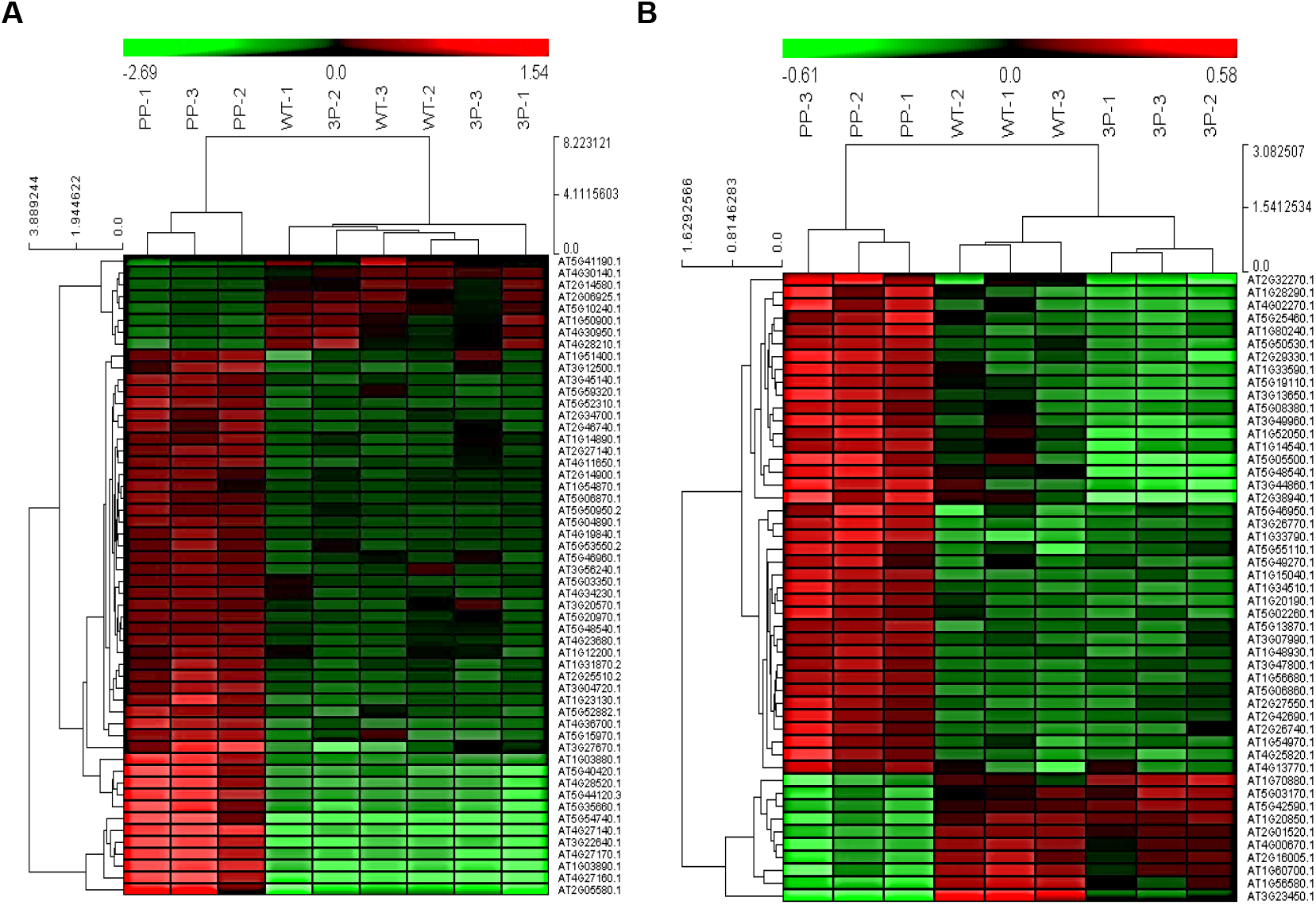
Comparative proteomics showing the functions of PDK1 and PEL1 in shaping the whole proteomes in *Arabidopsis*. Heat map displayed the abundance of 54 RCEs (restored CE proteins) in shoots (A) and 49 RCEs in roots (B) of wild type Col-0, *pdk1.1 pdk1.2* and *pdk1.1 pdk1.2 pel1*. “PP” refers to *pdk1.1 pdk1.2* double mutant and “3P” refers to *pdk1.1 pdk1.2 pel1* triple mutant. Three independent samples of WT (WT-1, 2, 3), *pdk1.1 pdk1.2* (PP-1, 2, 3) and *pdk1.1 pdk1.2 pel1* (3P-1, 2, 3) were collected and analyzed. Heat maps were generated using *log2*-transformed TMT values. Relative expression of the analyzed proteins was used to perform the hierarchical clustering analysis using Cluster3.0 (http://bonsai.hgc.jp/~mdehoon/software/cluster/software.htm) and Java Treeview software (http://jtreeview.sourceforge.net).

KEGG analysis of RCE proteins revealed the enriched metabolic pathways including lipids, carbohydrates, phenylpropanoid and amino acids, and involvement in multiple developmental processes and environmental adaptation (Table 1), which was consistent with the general growth defects of *pdk1.1 pdk1.2*. A nitrogen-regulated glutamine amidotransferase GAT 1_2.1 that represses shoot branching (*35*) increased in *pdk1.1 pdk1.2*, which is consistent with the solitary stem phenotype of *pdk1.1 pdk1.2*. Furthermore, increased DUF 642 family proteins DGR1 and DGR2 (*36*) and several root-hair-related proteins including SRPP (*37*), PRPL1 (*38*), GH9C1 (*39*), DER9 (*40*) and AGC2-1 (*41*) (a known PDK1 substrate) may account for the altered root development.

**Table 1.**
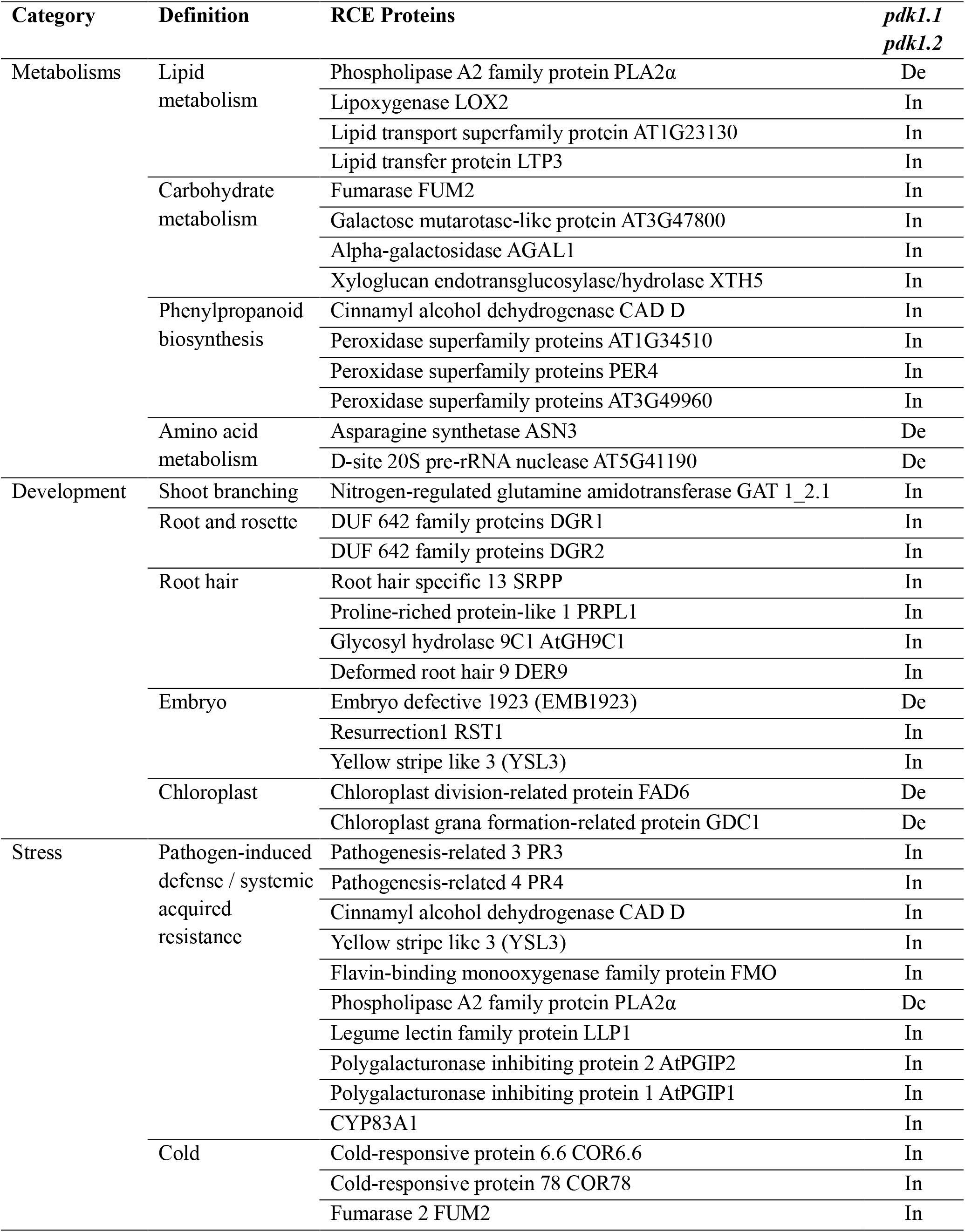
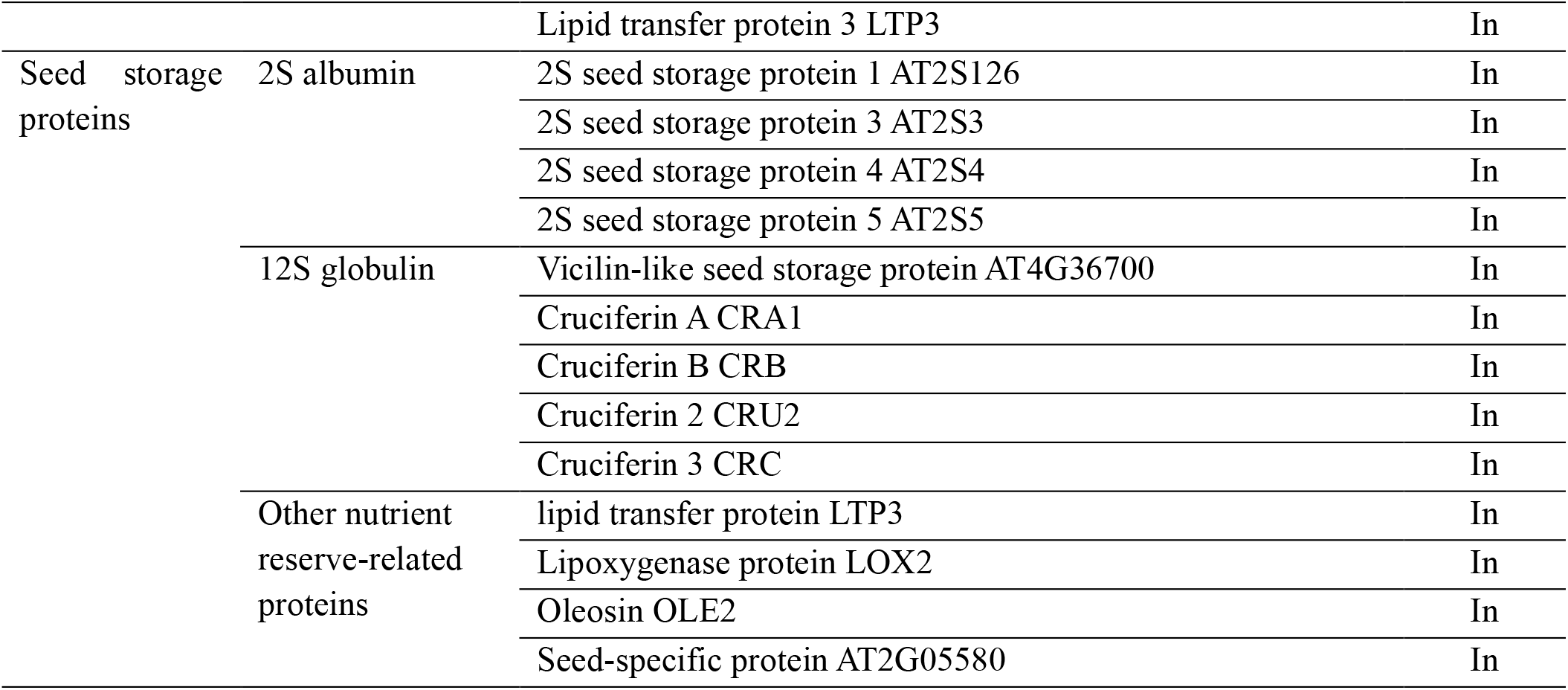
Identified RCEs (restored CE proteins) by analyzing Col-0, *pdk1*.*1 pdk1*.*2, pdk1*.*1 pdk1*.*2 pel1* mutants. Proteins were functionally categorized by KEGG pathway analysis and previous studies. In, increased; De, decreased.

A large number of pathogen-induced defense-related or systemic acquired resistance (SAR)-related proteins (*42*-*50*) accumulated in *pdk1.1 pdk1.2* (Table 1). Meanwhile, some abiotic stress-related proteins, especially cold acclimation/responsive proteins (*51*-*54*) significantly increased in *pdk1.1 pdk1.2* shoots. This is consistent with the previous studies showing that PDK1 positively regulates basal resistance in rice (*10*) and PDK1 is required for *P. indica*-induced growth promotion (*11*), and the rice PELOTA protein is involved in bacterial leaf blight resistance (*55*).

Notably, the TMT-based comparative proteomics analysis also showed that PEL1 and HBS1 presented unchanged protein abundance in *pdk1.1 pdk1.2* (Supplementary Fig. 10A, B). Next, we examined these known PDK1 substrates from the AGC family. Notably, there is an increase of D6PK protein, an essential downstream component of PDK1 (*12*), in *pdk1.1 pdk1.2*, but a relatively lower level of D6PK in *pdk1.1 pdk1.2 pel1* (Supplementary Fig. 11A). No dramatic changes were found for other detected AGC proteins in the proteomics data (Supplementary Fig. 11A, B). Therefore, it is very unlikely that changes of these AGC substrates might account for the rescue of *pdk1.1 pdk1.2* by the *pel1* mutation.

## Discussion

PDK1 is highly conserved in eukaryotes and is essential for growth and development of various organisms. *PDK1* deficiency results in severe growth defects or even lethality, which have impeded the studies on the underlying molecular mechanism, especially in mammals. Taking advantage of plant genetics and by screening for the suppressors that rescue the growth defects of T-DNA insertional *pdk1.1 pdk1.2* mutants, we here identify PEL1, which is a component of the PEL1-HBS1 mRNA surveillance complex and is essential for the mRNA quality control during protein translation. The mechanism for the *pel1* and *hbs1* mutations suppressing the *pdk1.1 pdk1.2* phenotype is their inability to degrade the aberrant mRNAs, leading to the production of truncated but functional PDK1 proteins (Fig. 5A, B). Our studies reveal that the PEL1-HBS1 complex coordinates the ribosome rescue and protein biosynthesis (Fig. 6A)

**Fig 6.**
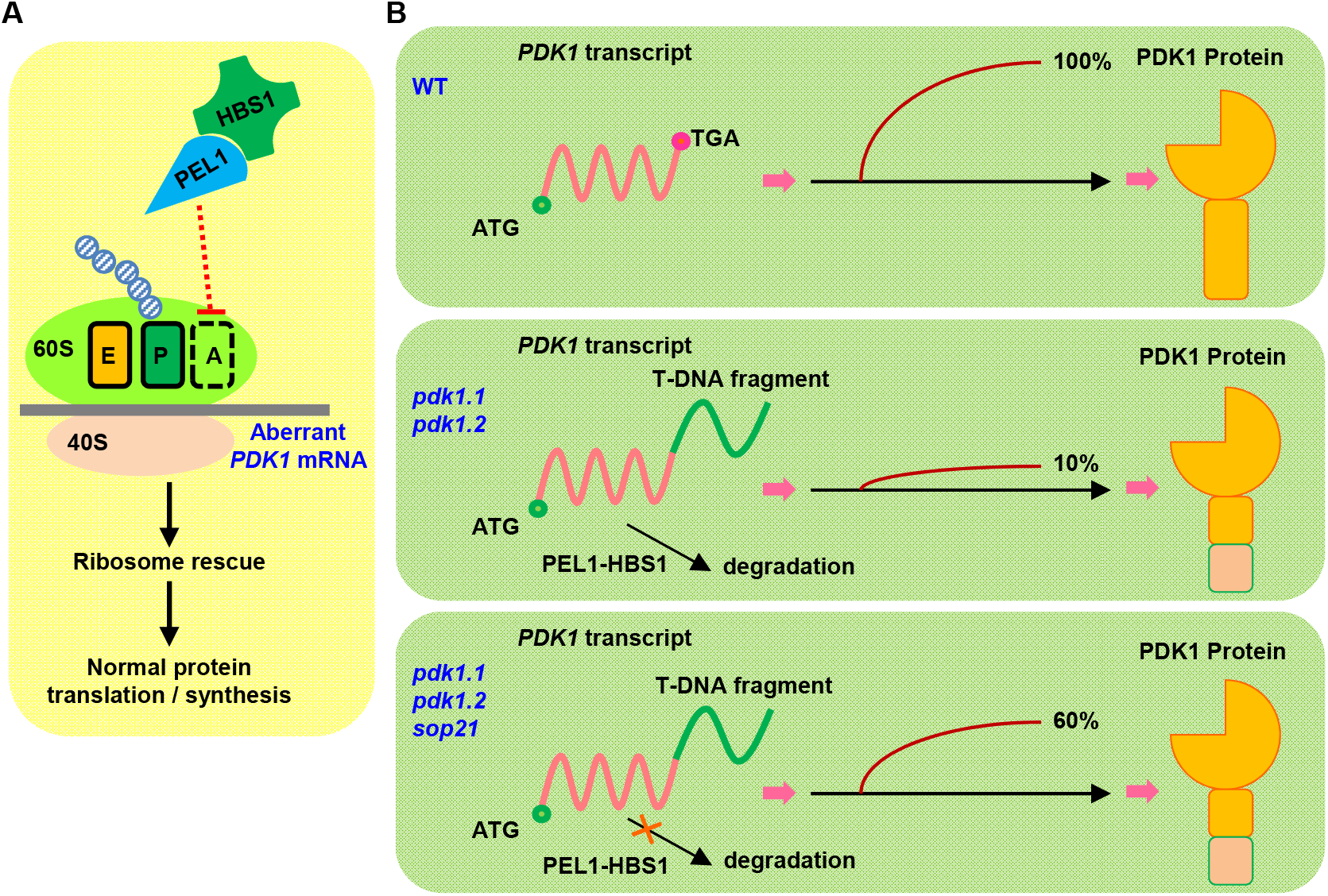
A proposed model showing the function of PEL1-HBS1 mRNA surveillance complex and how *sop21* suppresses *pdk1*.*1 pdk1*.*2* phenotypes. A. The PEL1-HBS1 complex regulates 80S ribosomes through translational surveillance to maintain the normal protein translation and plant growth. In the case of truncated *PDK1* transcripts in *pdk1.1 pdk1.2*, this complex could degrade these mRNAs without stop codon, thus promoting the recycling of stalled 80S ribosomes. A-site, ribosomal site most frequently occupied by aminoacyl-tRNA, which functions as acceptor for growing protein during peptide bond formation; P-site, ribosomal site most frequently occupied by peptidyl-tRNA, the tRNA carrying the chain of growing peptide; E-site, ribosomal site harbouring decylated tRNA on transit out from ribosome. **B**. A proposed model showing *sop21* mutation rescuing the defects of *pdk1.1 pdk1.2*: 1) In WT, the PDK1 transcripts have the stop codon, and it can be translated into 100% of PDK1 protein. 2) In the *pdk1.1 pdk1.2* T-DNA mutants, aberrant transcripts with fusion to partial T-DNA fragment will be recognized by the PEL1-HBS1 complex and thus get degraded, exhibiting PDK1 loss-of-function mutant defects. 3) The *sop21* mutations leads to the inefficient degradation of aberrant transcripts, which produce enough truncated PDK1 protein, maintaining normal growth of *pdk1.1 pdk1.2* plants.

PEL1-HBS1 complex plays important roles in the mRNA quality control. Our study is a typical example how this mRNA surveillance system regulates the stability of aberrant transcripts. Intriguingly, loss of function of DOM34 or PELOTA causes mitotic arrest in yeast and *Drosophila* and the *pelota* mutant mouse is lethal (*57*), whereas the *Arabidopsis* mutants *sop21, pel1*, and *hbs1* are viable, suggesting the different regulatory modes for protein translational regulation. Though the *pdk1.1 pdk1.2* double mutant exhibits pleiotropic defects throughout growth and development, the mutant can complete a life cycle (*12*). It is speculated that the difference might be related to the high postembryonic developmental plasticity in plants compared with animals, owing to the sessile life style during evolution.

Proteomics analysis indicates that the PEL1-HBS1 complex and PDK1 play pivotal roles in modulating the activity of the protein synthesis machinery under normal conditions. Notably, >80% RCE proteins are with increased abundances, and few decreased RCE proteins may due to the indirect/feedback regulation. The changed levels of most RCE proteins in *pdk1.1 pdk1.2* is reversed by the *pel1* mutation, suggesting a rescue at the whole proteome level. Given the lipid binding property of PDK1 and the localizations of PDK1 at PM, cytoplasm and certain ER-associated compartments, the pathway proposed here might be responsive to lipid dynamics at the membrane. Further characterization of the exact role of lipids in this process will help to elucidate the molecular mechanistic framework underlying the control of cell growth. In animals, PDK1 can phosphorylate one AGC kinase, AKT (aka Protein Kinase B, PKB) and S6K, to regulate protein biosynthesis through modulating 40S ribosomal protein 6S-A (RPS6A) and B (RPS6B), two subunits of ribosomes (*8, 34*). There are also two S6K homologues in *Arabidopsis*, previously reported to regulate protein synthesis via RPS6A/B (*58*). However, whether these two S6Ks are regulated by PDK1 requires further investigation.

*pdk1.1 pdk1.2* exhibits pleotropic growth and developmental defects, including significantly reduced fertility, which may be due to the changed levels of embryonic development-related proteins EMB 1923, RESURRECTION1 (RST1) (*32, 59*–*61*) and YELLOW STRIPE-LIKE3 (YSL3) (*62*). Notably, RST1 is a crucial regulator for RNA metabolism and thus the post-transcriptional gene silencing pathway (*32, 60*). Together with the previous biochemical evidence showing that RST1 forms a complex with HBS1 (SKI7) involved in post-transcriptional gene silencing (*32*), we speculated that the change of RST1 protein levels might be due to the altered status of HBS1 protein in *pdk1.1 pdk1.2*. We speculate that the increase level of RST1 might function as a compensation mechanism for the overall accelerated protein synthesis in *pdk1.1 pdk1.2*. Moreover, RST1 was recently identified as a regulator of the vacuolar protein degradation pathway (*61*), implying a role of PDK1 in the endomembrane trafficking process. Interestingly, precursors of two major storage proteins, 2S albumin and 12S globulin (*63, 64*) (Table 1) are significantly accumulated in *pdk1.1 pdk1.2* shoots. Likewise, a number of nutrient reserve-related proteins including lipid transfer protein LTP3 (*54*), lipoxygenase protein LOX2 (*65*), oleosin OLE2 (*66*) and seed-specific protein AT2G05580 (*67*) exhibited the same change in *pdk1.1 pdk1.2*. Storage proteins are actively synthesised at rough ER as precursor forms and then are transported into protein storage vacuole (PSV) during seed maturation (*68*). In higher plants, seed storage proteins are deposited in PSVs of dry seeds as a source of nitrogen for growth after seed germination (*68, 69*). Accumulation of seed storage proteins and nutrient reserve-related proteins in *pdk1.1 pdk1.2* shoots may result in the altered vegetative growth. This indicates that PDK1 represses seed storage proteins and nutrient reserve-related proteins in the vegetative tissues or the nitrogen utilization after seed germination.

It is noteworthy that the *pdk1.1 pdk1.2* phenotype is not fully rescued by *pel1* mutations, or by a truncated *PDK1.1N* transgene. We speculate that the truncated PDK1 protein only keeps partial functionality. Meanwhile, PDK1 may also regulate specific life activities through phosphorylating distinct substrates, including those well-characterised AGC kinases as well as possible others (*12, 18*). How these downstream pathways coordinate with each other, special-temporally, needs further investigation.

## Materials and Methods

### Materials and growth conditions

*Arabidopsis thaliana* lines used in this study were all in ecotype Columbia (Col-0) background. Seeds of Col-0 and various mutants, transgenic lines were germinated on MS (Murashige and Skoog, Duchefa) medium after two days’ stratification at 4C. Seedlings and plants were grown in a phytotron at 22°C with a 16-h light / 8-h dark photoperiod. Root growth measurements were performed using 14-day-old seedlings grown on MS.

Mutant lines *pdk1.1-2* (*pdk1.1*, SALK_113251C) (*12*), *pdk1.2-4* (*pdk1.2*, SALK_017433) (*12*), *pel1* (*pel1*, SALK_124403C), and *hbs1* (*hbs1*, CS857798) were obtained from ABRC (*70*) (Arabidopsis Biological Resource Centre) and were genotyped by using corresponding LB, RP and LP primers (Supplementary Table 2). *pPDK1.1::GUS* and *pPDK1.2::GUS* were reported previously (*12*). The floral dip method (*71*) was used for plant transformation.

### Reverse transcription-quantitative real-time RT-PCR (RT-qPCR) analysis

Total RNA was extracted from seedlings using TRIzolR reagent (Invitrogen), incubated with DNAase (TAKARA) and reverse transcribed (TAKARA). Transcription of corresponding genes and *ACTIN7* was analysed using SYBR Premix Ex Taq (TAKARA) with a BIO-RAD CFX Connect Real-Time System. Relative expression of examined genes was calculated by setting the gene expression level of wild type as “1” and was presented as average ± standard deviation (SD) from three independent biological replicates.

### Promoter::β-glucuronidase (GUS) staining for expression pattern analysis

*pPDK1.1::GUS* and *pPDK1.2::GUS* transgenic lines were reported previously (*12*), and *pPEL1::GUS* and *pHBS1::GUS* lines were cloned with a modified pCambia1300 binary vector (*72*) using primers listed in Supplementary Table 2. Stable transgenic lines were stained at 37°C for 1 h, in GUS solution [0.5 mg/mL 5-bromo-4-chloro-3-indolyl-β-d-glucuronic acid (X-Gluc), 0.5 mM potassium ferricyanide (K_4_[Fe(CN)_6_]·3H_2_O), 0.5 mM potassium ferrocyanide (K_3_[Fe(CN)_6_]), 0.1% (v/v) Triton X-100, 10 mM ethylenediaminetetraacetic acid (EDTA) and 0.1 M sodium phosphate (NaH_2_PO_4_); pH 7.0] (*12*). Three independent lines were analysed in detail for different tissues and stages, and they all showed similar expression patterns. Samples were imaged by a stereomicroscope (Nikon SMZ1500).

### Yeast two-hybrid (Y2H) assay

Y2H assays were performed as reported previously (*73*). Coding sequences of *PEL1, PEL1*^*T43A*^ and *PEL1*^*T43E*^ were amplified by PCR with PEL1-F (*BamH*I)-3 and PEL1-R (*BamH*I) primers, and were then subcloned into the pGBKT7 vector (Clontech). Coding sequences of HBS1 [using primers HBS1-F (*EcoR*I) and HBS1-R (*BamH*I)] and HBS1 [using primers HBS1-2-F (*Nde*I) and HBS1-2-R (*BamH*I)] were amplified and subcloned into pGADT7 vectors (Clontech). Bait and prey plasmids were co-transformed into the yeast strain AH109 according to the manufacture’s introduction (Clontech). Transformants were selected on SD (-Leu/-Trp) solid medium. For auxotroph assays, four individual colonies were cultured in liquid SD (-Leu/-Trp) medium overnight, and approximately 10 µL of each sample at different dilutions (as indicated in the figure legends) was dropped on SD (-Leu/-Trp/-His) medium supplemented with 0.5 mg/mL X-α-Gal or on SD (-Leu/-Trp/-His/-Ade) medium, respectively, with 1 mM 3-amino-1,2,4-triazole (3-AT), and grown at 30°C for 3 days. Colonies showing continuous growth with a blue colour represented interactions.

### Bimolecular fluorescence complementation (BiFC) assay

For BiFC assay, cDNAs encoding *PDK1.1, PDK1.2, PEL1* and *HBS1* were cloned into the pENTR plasmid with BP reactions. Afterwards, LR reactions were conducted with the 35S::GW-nYFP and 35S::GW-nYFP destination vectors (*74*), resulting in *35S::PDK1.1-nYFP, 35S::PDK1.1-cYFP, 35S::PDK1.2-nYFP, 35S::PDK1.2-cYFP, 35S::PEL1-nYFP* and *35S::HBS1-cYFP*, respectively. Resultant constructs with control blank vectors were co-expressed in *N. benthamiana* leaves and yellow fluorescence was observed by a Leica SP8 confocal laser scanning microscope, using an argon laser excitation wavelength of 488 nm after infiltration for 48 days.

### Subcellular localization and co-localization studies

For subcellular localization studies, cDNAs encoding *PDK1.1, PDK1.2, PEL1* and *HBS1* were first cloned into the pENTR plasmid with BP reactions. Afterwards, LR reactions were conducted with the pGWB605 destination vector, resulting in *pGWB605-35S::PDK1.1-GFP, pGWB605-35S::PDK1.2-GFP, pGWB605-35S::PEL1-GFP* and *pGWB605-35S::HBS1-GFP*, respectively. PDK1.1-GFP, PDK1.2-GFP, PEL1-GFP, HBS1-GFP and ER-mCherry (*75*) fusion proteins were transiently expressed in *N. benthamiana* leaves (*76*). For mCherrry fusion studies, *PDK1.1* and *PEL1* were cloned into the pB7m24GW2 destination vector. *35S::GFP-HDEL* was used for the ER reporter. The infiltrated leaves were harvested 2 days after infiltration and observed using an Olympus confocal microscope (Olympus, FV10i). *PDK1.1-YFP, PDK1.2-YFP, PEL1-YFP*, and *HBS1-YFP* were cloned into the pA7 plasmid and transiently expressed in leaf protoplasts of wild type, *Arabidopsis* seedlings expressing ER-mCherry (*75*), or PIP2-RFP (*77*). Transformed protoplasts were harvested 12 hours after transformation and observed using an Olympus confocal microscope (Olympus, FV10i).

For *35S::PDK1.1-GFP, 35S::PDK1.2-GFP* and *35S::PEL1-GFP* transgenic plants, entry vectors were reacted with pB7FWG2,0 plasmids for GFP fusion expression. Transformation was performed with the floral dip method (*71*) with the *Agrobacteria* stain GV3101.

Images were captured with following excitation (Ex) and emission (Em) wavelengths (Ex/Em): GFP 488 nm/501-528 nm; /YFP 490 nm/520-550 nm; mCherry/RFP 543 nm/620-630 nm; DAPI 405 nm/437-476 nm.

### Protein extraction and Western blot analysis

To examine the protein levels of FLAG- and GFP-tagged proteins, approximately 100 mg of plant tissues were frozen in liquid nitrogen, ground thoroughly, and homogenized in 100 μL protein extraction buffer [20 mM Tris-HCl, pH 7.5, 150 mM NaCl, 0.5% (v/v) Tween-20, 1 mM EDTA, 1 mM DTT] containing a protease inhibitor cocktail (cOmplete, Roche) and a protein phosphatase inhibitor tablet (PhosSTOP, Roche). After addition of SDS loading buffer, the samples were heated at 65°C for 5 min, resolved by 10% (v/v) SDS-PAGE and transferred to PVDF membranes. FLAG-tagged proteins were detected by a mouse anti-FLAG antibody (M20008, 1:2,000, Abmart). GFP-tagged proteins were detected with a mouse anti-GFP antibody (M20004, 1:2,000, Abmart) or a mouse anti-GFP HRP-conjugated antibody (130-091-833, 1:2,000, MACS Molecular). His-tagged proteins were detected by a mouse anti-His antibody (sc-8036, 1:3000, Santa Cruz Biotechnology). GST-tagged proteins were detected by a mouse anti-GST antibody (sc-138, 1:3000, Santa Cruz Biotechnology). Actin was detected by a mouse anti-actin antibody (M20009, 1:2,000, Abmart). HRP activity was detected by the Supersignal Western Detection Reagents (Thermo Scientific). After incubated with a primary mouse antibody, the PVDF membrane was then incubated with a goat anti-mouse immunoglobulin G AP-conjugated secondary antibody (ab97020, 1:5000, Abcam). AP activity was detected by BCIP/NBT kit (Invitrogen) according to the supplier’s instructions.

### Protein expression and *in vitro* kinase assay

Coding regions of PDK1.1, PEL1 and HBS1 were amplified with corresponding primers, and subcloned into vectors pET28a (Novagen) or pGEX-4T-1 (GE Healthcare) respectively. Proteins were recombinantly expressed in *Escherichia coli* (strain BL21) by supplementing with 1 mM or 0.2 mM isopropyl-β-D-thiogalactopyranoside (IPTG, induced at either 28°C for 3 h or 16°C for 16 h). Fusion proteins with His tag were purified using Ni-NTA His binding resin (Novagen) and those with GST tag was purified by glutathione sepharose (Novagen).

Kinase activity assay was performed according to previous reports(*12, 22, 78*) with minor modifications. Assay was initiated by adding 1 µg recombinant His-PDK1.1 in a total volume of 40 µL containing 50 mM Tris-HCl, pH 7.5, 5 mM MgCl_2_, 2 mM CaCl_2_, 1 mM DTT (1,4-dithiothreitol), 0.1 mM ATP (Adenosine 5’-triphosphate), 5 μCi [γ-^32^P]ATP (NEC902A; Perkin-Elmer), and 10 µg of substrate (recombinant His-PEL1, His-PEL1^T43A^, or GST-HBS1). Reactions were incubated at 30°C for 45 min and terminated by adding 2× SDS loading buffer. After boiling for 5 min, the reaction products were fractionated by SDS-PAGE (sodium dodecyl sulfate-polyacrylamide gel electrophoresis), and the radioactivity was collected by a phosphor screen. After 10 hour, the phosphor screen was imaged by autoradiography (Fujifilm FLA 9000 plus DAGE).

### A tandem mass tag (TMT)-based comparative proteomics analysis

A tandem mass tag (TMT)-based comparative proteomics analysis is performed according to Thompson’s research (*79*). Wild type, *pdk1.1 pdk1.2, pdk1.1 pdk1.2 pel1* seedlings grow for 14 days on the 1/2MS dishes. One gram shoots (aerial parts) and 0.6 g roots (underground parts) of three genotypes of seedlings were set as group1 and group2 respectively. Experiments were biological repeated for three times.

Samples were frozen in liquid nitrogen and ground homogeneously. 5 times volume of TCA/acetone (1:9) was added to the powder and mixed by vortexing. The mixture was placed at −20°C for 4 h, and centrifuged at 6, 000 g for 40 min at 4°C. The supernatant was discarded. The pre-cooling acetone was added to wash for three times. The pellet was air dried. 30 times volume of SDT buffer was added to 20-30 mg powder, mixed and boiled for 5 min. The lysate was sonicated and then boiled for 15 min. After centrifuged at 14, 000 g for 40 min, the supernatant was filtered with 0.22 µm filters. The filtrate was quantified with the BCA Protein Assay Kit (Bio-Rad, USA). The sample was stored at −80°C.

20 µg of proteins for each sample were mixed with 5× loading buffer respectively and boiled for 5 min. Proteins were separated with 12.5% SDS-PAGE (constant current 14 mA, 90 min) and bands were visualized by Coomassie Blue R-250 staining.

200 μg of proteins for each sample were incorporated into 30 μl SDT buffer (4% SDS, 100 mM DTT, 150 mM Tris-HCl, pH 8.0). The detergent, DTT and other low-molecular-weight components were removed using UA buffer (8 M Urea, 150 mM Tris-HCl pH 8.0) by repeated ultrafiltration (Microcon units, 10 kDa). Then 100 μl iodoacetamide (100 mM IAA in UA buffer) was added to block reduced cysteine residues and the samples were incubated for 30 min in darkness. The filters were washed with 100 μl UA buffer three times and then 100 μl 100 mM TEAB buffer twice. Finally, the protein suspensions were digested with 4 μg trypsin (Promega) in 40 μl TEAB buffer overnight at 37°C, and the resulting peptides were collected as a filtrate. The peptide content was estimated by UV light spectral density at 280 nm using an extinctions coefficient of 1.1 of 0.1% (w/v) solution that was calculated on the basis of the frequency of tryptophan and tyrosine in vertebrate proteins.

100 μg peptide mixture of each sample was labelled using TMT reagent according to the manufacturer’s instructions (Thermo Fisher Scientific) and analysed on an Orbitrap Fusion Lumos (Thermo Scientific) mass spectrometer coupled with Ultimate 3000 RSLC nano system. 4 μl of each fraction was injected for nano LC-MS/MS analysis. The peptide mixture (1 μg) was loaded onto the Acclaim PepMap 100 analytical column (75 μm × 15 cm, C18, 3 μm, Thermo Scientific) in buffer A (0.1% Formic acid) and separated with a linear gradient of buffer B (80% acetonitrile and 0.1% Formic acid) at a flow rate of 300 nl/min. The electrospray voltage of 2.1 kV versus the inlet of the mass spectrometer was used. Mass spectrometer was operated in the data-dependent mode to switch automatically between MS and MS/MS acquisition with a cycle time of 3 second. Survey full-scan MS spectra (m/z 375-1800) were acquired with a mass resolution of 120K, followed by sequential high energy collisional dissociation (HCD) MS/MS scans with a resolution of 50K. In all cases, one microscan was recorded using dynamic exclusion of 40 seconds. For MS/MS, precursor ions were activated using 38% normalized collision energy.

MS/MS spectra were analysed using ProteinDiscoverer− Software 2.1 against TAIR10_pep_20101214 database and decoy database with following parameters. The highest score for a given peptide mass (best match to that predicted in the database) was used to identify parent proteins. Parameters for protein searching were set as follows: tryptic digestion with at most two missed cleavages, carbamidomethylation of cysteines as fixed modification, and oxidation of methionines and protein N-terminal acetylation as variable modifications. Peptide spectral matches were validated based on q values at a 1% false discovery rate (FDR).

Proteins were considered differentially expressed when they displayed significant changes (more than 1.2-fold and Student’s *t* test, *P* value < 0.05).

The FASTA protein sequences of differentially changed proteins were blasted against the online Kyoto Encyclopedia of Genes and Genomes (KEGG) database (http://geneontology.org/) to retrieve their KOs and were subsequently mapped to pathways in KEGG. The corresponding KEGG pathways were extracted.

### Quantification and statistics

Lateral root numbers were counted directly. For measurements of silique length, primary root length and leaf area, photos were analysed with the Image J program (https://imagej.nih.gov/ij/download.html). Fluorescence intensity of reporter lines was analysed and quantified by Fiji (https://fiji.sc/) (*80*). Data visualisation and statistics were performed with GraphPad Prism8. Student’s t-test was used for comparing two data sets, and one-way ANOVA was performed for multiple comparisons.

## Acknowledgements

We gratefully acknowledge the *Arabidopsis* Biological Resource Centre (ABRC) for providing T-DNA insertional mutants, and Prof. Remko Offringa for sharing published seeds. We thank Xixi Zhang for the pDONR-P4P1r-mCherry plasmid, Alexander Johnson and Hana Semeradova for helpful comments. The study was supported by National Science Foundation of China (NSFC, 31721001, to H.-W. X.), “Ten-Thousand Talent Program” (to H.-W. X.) and Collaborative Innovation Center of Crop Stress Biology, Henan Province, and Austrian Science Fund (FWF): I 3630-B25 (to J. F.). S.T. was funded by a European Molecular Biology Organization (EMBO) long-term postdoctoral fellowship (ALTF 723-2015).

## Author contributions

W. K. performed acquisition of most of the data on *sop21* as well as analysis and interpretation of data and drafted the article. S. T. generated and analysed various mutant materials, as well as the generation of *pdk1.1 pdk1.2* EMS population and marker lines, suppressor screening and subcellular localization analysis. Q. Z. performed the backcross and the NGS analysis. D.-L. L. performed RT-qPCR and semi-quantitative analysis for *pel1* crosses. H.-W. X. is responsible for conception and design. Z.-H. X. and J.F. helped design experiments. W.K., S.T., J.F. and H.-W. X. wrote the manuscript, and all authors revised and approved it.

## Conflict of Interests

The authors declare no competing financial interests.

## Data and materials availability

All data and materials necessary to evaluate the conclusions in paper or supplementary materials are available.

## Supplementary Information

Supplementary Figures. 1 to 11

Supplementary Tables. 1 to 2

## Legends of Supplementary figures

**Supplementary Fig. 1 Expression profiles of *PDK1***.***1* and *PDK1***.***2***.

A. Reverse transcription-quantitative PCR (RT-qPCR) analysis reveals the expression of *PDK1.1* and *PDK1.2* in various tissues. *ACTIN7* gene was amplified and used as an internal control. Experiments were biologically repeated 3 times and data are presented as means ± SD (standard derivation). n = 3. Different letters represent significant difference, *P <* 0.05, by one-way ANOVA with a Tukey multiple comparison test.

B-M. Promoter-reporter gene (*GUS*) fusion studies show the expression of *PDK1.1* in 7-day-old seedlings (B) including shoot apical meristem (SAM, C), cotyledon (D), primary root tip (E), lateral root (J) and floral tissues (mainly in pollens, L); and similar expression pattern of *PDK1.2* (F-I, K, M). Scale bars, 2 mm (B, F), 500 μm (D, H, L, M), 200 μm (C, G, E, I) or 100 μm (J, K). Transgenic lines were confirmed and three independent lines were analyzed. Representative images are shown.

**Supplementary Fig. 2 Identification of the *pel1* mutant**.

A. Schematic representation of *PEL1* gene and position of T-DNA insertion. Introns, exons and non-coding regions are indicated by lines, black or blank boxes. Positions of primers are indicated.

B. Identification of homozygous *pel1* mutant. Genomic DNAs were used as template for PCR amplification and homozygous lines presents a single amplified fragment when using LBa1/PEL1-RP primers.

C. RT-qPCR analysis confirmed the deficient *PEL1* expression in *pel1*. Total RNA of 7-day-old Col-0 and *pel1* seedlings was extracted and used for analysis. *ACTIN7* gene was amplified and used as an internal control. Experiments were biologically repeated 3 times and data are presented as means ± SD. n = 3. *P* value was calculated by an unpaired Student’s t-test.

D. Quantification of the rosette area revealed that *pel1* mutations partially rescued the phenotype of *pdk1.1 pdk1.2*. n = 16, 21, 15, 16, and 12, respectively. Different letters represent significant difference, *P <* 0.05, by one-way analysis of variance (ANOVA) with a Tukey multiple comparison test.

E-F. A representative photo showed that the *pel1* mutations partially rescued the inflorescence phenotype of *pdk1.1 pdk1.2*. Scale bars, 2 cm.

**Supplementary Fig. 3 Expression profiles of *PEL1***.

A. RT-qPCR analysis revealed the *PEL1* expression in various tissues. *Actin7* gene was amplified and used as an internal control. Experiments were biologically repeated 3 times and data are presented as means ± SD. n = 3. Different letters represent significant difference, *P <* 0.05, by one-way ANOVA with a Tukey multiple comparison test.

B-G. Promoter-reporter gene (GUS) fusion studies reveals the *PEL1* expression in 7-day-old seedlings (B) including primary roots (C), shoots (D) and lateral roots (E), 21-day-old seedlings (F), and floral organs (G). Transgenic lines were confirmed and three independent lines were analyzed. Scale bars, 2 mm (B), 200 μm (C), 500 μm (D, G), 100 μm (E), or 1 cm (F). Representative images are shown.

**Supplementary Fig. 4 Loss of function of *PEL1* suppresses the defective growth of *pdk1***.***1 pdk1***.***2*, confirmed by complementation**.

A. Western blot analysis confirms the *PEL1-FLAG* expression in *pdk1.1 pdk1.2 sop21* transgenic lines. An anti-FLAG (upper panel) and an anti-Actin (bottom panel) antibody was used respectively.

B. Expression of PEL1 protein in *sop21* plants restored the phenotype of *pdk1.1 pdk1.2*. Five-(top) or three-(bottom) week-old Col-0, *pdk1.1 pdk1.2, pdk1.1 pdk1.2 sop21* and *35S::PEL1-FLAG* (in *pdk1.1 pdk1.2 sop21*) plants were observed and shown. Scale bars, 2 cm.

C. Western blot analysis confirmed the PEL1-FLAG protein expression in Col-0 background. An anti-FLAG (upper panel) and an anti-Actin (bottom panel) antibody was used respectively.

D. Overexpression of *PEL1* did not show any obvious phenotypes. Five-(top) or three-(bottom) week-old Col-0 and *35S::PEL1-FLAG* (in Col-0) plants were observed and representative photos are shown. Scale bars, 2 cm.

**Supplementary Fig. 5 Sequence alignment among *Arabidopsis thaliana* HBS1, *Homo sapiens* HBS1 and *Saccharomyces cerevisiae* HBS1**. Protein sequences were obtained from NCBI, including AtHBS1 (AED91575.1), HsHBS1 (NP_006611.1) and ScHBS1 (CAA82163.1). Alignment was performed using the DNAMAN Software with default settings. The three orthologues show 38.60% identity. The similarity was shown in different colours: back, 100% identity; grey, ≥75%.

**Supplementary Fig. 6 Identification of *hbs1* mutants**.

A. RT-qPCR analysis reveals the *HBS1* expression in various tissues. *Actin7* gene was amplified and used as an internal control. Experiments were biologically repeated 3 times and data are presented as means ± SD. n = 3. Different letters represent significant difference, *P <* 0.05, by one-way ANOVA with a Tukey multiple comparison test.

B. Schematic representation of *HBS1* gene and position of T-DNA insertion. Introns, exons and non-coding regions are indicated by lines, black or blank boxes. Positions of primers are indicated.

C. Identification of homozygous *hbs1* mutant. Genomic DNA was used as template for PCR amplification and homozygous lines presents a single amplified fragment when using P745/HBS1-LP primers.

D. RT-qPCR analysis confirmed the significantly reduced *HBS1* expression in *hbs1* mutant. Total RNAs of 7-day-old WT and *hbs1* seedlings were extracted and used for analysis. *ACTIN7* gene was amplified and used as an internal control. Experiments were biologically repeated 3 times and data are presented as means ± SD. *P* value was calculated by an unpaired Student’s t-test.

**Supplementary Fig. 7 Phenotype of multiple combinations of *pdk1***.***1 pdk1***.***2* double mutants**.

A-B. A schematic picture showing the positions of T-DNA insertions in various alleles of *pdk1.1* and *pdk1.2*. The primers used for genotyping *pdk1.1* (*pdk1.2-2*) and *pdk1.2* (*pdk1.2-4*) and RT-qPCR analysis are also indicated with arrows.

C. A representative photo showing the phenotype of different combinations of *pdk1.1 pdk1.2* double mutants. 25 days old. Scale bar, 2 cm.

**Supplementary Fig. 8 Increased expression of PDK1 N-terminal fragments rescued the growth defects of *pdk1***.***1 pdk1***.***2***.

A. Semi-quantitative RT-PCR analysis with primers targeting the N-terminal fragments (beforeT-DNA insertions) revealed that N-terminal fragments of *PDK1.1* and *PDK1.2* transcript (PDK1.1N and PDK1.2N) exhibited increased levels in *pdk1.1 pdk1.2 sop21, pdk1.1 pdk1.2 pel1* and *pdk1.1 pdk1.2 hbs1*, respectively, compared to that in *pdk1.1 pdk1.2. ACTIN7* gene was amplified and used as an internal control (bottom).

B. CaMV 35S-driven overexpression of PDK1 N-terminal fragment rescued the growth defects of *pdk1.1 pdk1.2*. A representative photo of 20-day-old Col-0, *pdk1.1 pdk1.2, 35S::Venus-PDK1.1, 35S::Venus-PDK1.2, 35S::Venus-PDK1.1N, 35S::Venus-PDK1.2N, 35S::Venus-PDK1.1C* and *35S::Venus-PDK1.2C* plants (all in *pdk1.1 pdk1.2* background) grown in soil are shown. Scale bar, 2 cm.

**Supplementary Fig. 9 Subcellular localization of PDK1**.**1-YFP, PDK1**.**2-YFP, PEL1-YFP, and HBS1-YFP**.

A-C. Fusion proteins PDK1.1-YFP, PDK1.2-YFP, PEL1-YFP, and HBS1-YFP were transiently expressed in *Arabidopsis* leaf protoplasts and fluorescence were observed (a). Endoplasmic reticulum-specific ER-mCherry (a), plasma membrane-specific PIP2-RFP (b), and nuclear-specific dye DAPI (c) were used to confirm the location at plasma membrane, endoplasmic reticulum, or nucleus. Scale bars, 10 µm.

D. Western blot revealing the integrity of PDK1.1-GFP protein in *35S::PDK1.1-GFP* transgenic plants. Upper panel, anti-GFP antibody; bottom, Ponceau stain.

E. Fluorescence observations show the endoplasmic reticulum localization of PDK1.1, PDK1.2, PEL1 and HBS1. Fusion proteins PDK1.1-GFP, PDK1.2-GFP, PEL1-GFP and HBS1-GFP were transiently expressed with ER-specific ER-mCherry proteins in tobacco leaves. Scale bars, 50 μm.

**Supplementary Fig. 10 PDK1 doesn’t affect the amounts of PEL1 and HBS1 proteins**. Abundance of PEL1 and HBS1 proteins is not changed in roots of 14-day-old *pdk1.1 pdk1.2* seedlings by TMT-based comparative proteomics analysis. “PP” refers to *pdk1.1 pdk1.2* double mutant and three independent samples of WT (WT-1, WT-2, WT-3) and *pdk1.1 pdk1.2* (PP-1, PP-2, PP-3) were collected and analyzed. Heat maps were generated using *log2*-transformed TMT values. Relative amount of PEL1 and HBS1 proteins was used to perform the hierarchical clustering analysis using Cluster3.0 (http://bonsai.hgc.jp/~mdehoon/software/cluster/software.htm) and Java Treeview software (http://jtreeview.sourceforge.net). Euclidean distance algorithm for similarity measure and average linkage clustering algorithm (clustering uses the centroids of observations) for clustering were selected when performing hierarchical clustering.

**Supplementary Fig. 11 Abundance of AGC protein kinases in *pdk1***.***1 pdk1***.***2* or *pdk1***.***1 pdk1***.***2 pel1***.

Abundance of the AGC family of proteins detected in shoots (A) or roots (B) of 14-day-old *pdk1.1 pdk1.2* and *pdk1.1 pdk1.2 pel1* seedlings by TMT-based comparative proteomics analysis. Data are presented as means ± SD. n = 3. Different letters represent significant difference, *P <* 0.05, by one-way ANOVA with a Tukey multiple comparison test.

**Table S1** | **Candidate HBS1 proteins in *Arabidopsis thaliana* by homologous analysis using *Saccharomyces cerevisiae* HBS1 (ScHBS1) and *Homo sapiens* HBS1 (HsHBS1L)**. Top five *Arabidopsis* homologs of ScHBS1 and HsHBS1L are shown and corresponding properties were obtained through the TAIR website (http://www.arabidopsis.org/).

**Table S2** | **Primers used in this study**. Added restriction enzymes are indicated and underlined.

## References

1. T. Tsuboi, K. Kuroha, K. Kudo, S. Makino, E. Inoue, I. Kashima, T. Inada, Dom34:hbs1 plays a general role in quality-control systems by dissociation of a stalled ribosome at the 3’ end of aberrant mRNA. Mol. Cell. 46, 518–529 (2012).

2. C. J. Shoemaker, D. E. Eyler, R. Green, Dom34:Hbs1 promotes subunit dissociation and peptidyl-tRNA drop-off to initiate no-go decay. Science. 330, 369–372 (2010).

3. D. R. Alessi, Discovery of PDK1, one of the missing links in insulin signal transduction. Biochem. Soc. Trans. 29, 1–14 (2001).

4. A. Mora, D. Komander, D. M. F. Van Aalten, D. R. Alessi, PDK1, the master regulator of AGC kinase signal transduction. Semin. Cell Dev. Biol. 15, 161–170 (2004).

5. A. Storz, P., Toker, 3 -phosphoinositide-dependent kinase-1 PDK-1 in PI 3-kinase signaling. Front. Biosci. 7, 886–902 (2002).

6. A. Casamayor, P. D. Torrance, T. Kobayashi, J. Thorner, D. R. Alessi, Functional counterparts of mammalian protein kinases PDK1 and SGK in budding yeast. Curr. Biol. 9 (1999), pp. 186–197.

7. M. Inagaki, T. Schmelzle, K. Yamaguchi, K. Irie, M. N. Hall, K. Matsumoto, PDK1 homologs activate the Pkc1-mitogen-activated protein kinase pathway in yeast. Mol. Cell. Biol. 19, 8344–52 (1999).

8. F. Rintelen, H. Stocker, G. Thomas, E. Hafen, PDK1 regulates growth through Akt and S6K in Drosophila. Proc. Natl. Acad. Sci. 98, 15020–15025 (2001).

9. M. A. Lawlor, A. Mora, P. R. Ashby, M. R. Williams, V. Murray-Tait, L. Malone, A. R. Prescott, J. M. Lucocq, D. R. Alessi, Essential role of PDK1 in regulating cell size and development in mice. EMBO J. 21, 3728–3738 (2002).

10. H. Matsui, A. Miyao, A. Takahashi, H. Hirochika, Pdk1 kinase regulates basal disease resistance through the OsOxi1-OsPti1a phosphorylation cascade in rice. Plant Cell Physiol. 51, 2082–2091 (2010).

11. I. Camehl, C. Drzewiecki, J. Vadassery, B. Shahollari, I. Sherameti, C. Forzani, T. Munnik, H. Hirt, R. Oelmüller, The OXI1 kinase pathway mediates Piriformospora indica-induced growth promotion in Arabidopsis. PLoS Pathog. 7, e1002051 (2011).

12. S. Tan, X. Zhang, W. Kong, X.-L. Yang, G. Molnár, Z. Vondráková, R. Filepová, J. Petrášek, J. Friml, H.-W. Xue, The lipid code-dependent phosphoswitch PDK1–D6PK activates PIN-mediated auxin efflux in Arabidopsis. Nat. Plants. 6, 556–569 (2020).

13. Y. Xiao, R. Offringa, PDK1 regulates auxin transport and Arabidopsis vascular development through AGC1 kinase PAX. Nat. Plants. 6, 544–555 (2020).

14. A. C. N. Dittrich, T. P. Devarenne, Characterization of a PDK1 Homologue from the Moss Physcomitrella patens. Plant Physiol. 158 (2012), pp. 1018–1033.

15. S. Tan, C. Luschnig, J. Friml, Pho-view of Auxin: Reversible Protein Phosphorylation in Auxin Biosynthesis, Transport and Signalling. Mol. Plant. 14, 151–165 (2021).

16. R. G. Anthony, R. Henriques, A. Helfer, T. Mészáros, G. Rios, Testerink, T.C. Munnik, M. Deák, C. Koncz, L. Bögre, A protein kinase target of a PDK1 signalling pathway is involved in root hair growth in Arabidopsis. EMBO J. 23, 572–581 (2004).

17. M. Deak, A. Casamayor, R. A. Currie, C. Peter Downes, D. R. Alessi, Characterisation of a plant 3-phosphoinositide-dependent protein kinase-1 homologue which contains a pleckstrin homology domain. FEBS Lett. 451, 220–226 (1999).

18. H. Zegzouti, W. Li, T. C. Lorenz, M. Xie, C. T. Payne, K. Smith, S. Glenny, G. S. Payne, S. K. Christensen, Structural and functional insights into the regulation of Arabidopsis AGC VIIIa kinases. J. Biol. Chem. 281, 35520–35530 (2006).

19. M. C. Rentel, D. Lecourieux, F. Ouaked, S. L. Usher, L. Petersen, H. Okamoto, H. Knight, S. C. Peck, C. S. Grierson, H. Hirt, M. R. Knight, OXI1 kinase is necessary for oxidative burst-mediated signalling in Arabidopsis. Nature. 427, 858–861 (2004).

20. R. G. Anthony, R. Henriques, A. Helfer, T. Mészáros, G. Rios, C. Testerink, T. Munnik, M. Deák, C. Koncz, L. Bögre, A protein kinase target of a PDK1 signalling pathway is involved in root hair growth in Arabidopsis. EMBO J. 23, 572–581 (2004).

21. J. Friml, X. Yang, M. Michniewicz, D. Weijers, A. Quint, O. Tietz, R. Benjamins, P. B. F. Ouwerkerk, K. Ljung, G. Sandberg, P. J. J. Hooykaas, K. Palme, R. Offringa, A PINOID-dependent binary switch in apical-basal PIN polar targeting directs auxin efflux. Science. 306, 862–865 (2004).

22. H. Zegzouti, R. G. Anthony, N. Jahchan, L. Bogre, S. K. Christensen, Phosphorylation and activation of PINOID by the phospholipid signaling kinase 3-phosphoinositide-dependent protein kinase 1 (PDK1) in Arabidopsis. Proc. Natl. Acad. Sci. 103, 6404–6409 (2006).

23. M. Zourelidou, I. Muller, B. C. Willige, C. Nill, Y. Jikumaru, H. Li, C. Schwechheimer, The polarly localized D6 PROTEIN KINASE is required for efficient auxin transport in Arabidopsis thaliana. Development. 136, 627–636 (2009).

24. P. Marhava, A. E. L. Bassukas, M. Zourelidou, M. Kolb, B. Moret, A. Fastner, W. X. Schulze, P. Cattaneo, U. Z. Hammes, C. Schwechheimer, C. S. Hardtke, A molecular rheostat adjusts auxin flux to promote root protophloem differentiation. Nature. 558, 297–300 (2018).

25. Y. Zhang, J. He, S. McCormick, Two Arabidopsis AGC kinases are critical for the polarized growth of pollen tubes. Plant J. 58, 474–484 (2009).

26. D. R. Page, U. Grossniklaus, The art and design of genetic screens: Arabidopsis thaliana. Nat. Rev. Genet. 3, 124–136 (2002).

27. R. S. Allen, K. Nakasugi, R. L. Doran, A. A. Millar, P. M. Waterhouse, Facile mutant identification via a single parental backcross method and application of whole genome sequencing based mapping pipelines. Front. Plant Sci. 4, 362 (2013).

28. T. Csorba, A. Auber, A. Schamberger, The nonstop decay and the RNA silencing systems operate cooperatively in plants. Nucleic Acids Res. 46, 4632–4648 (2018).

29. P. Qin, S. Fan, L. Deng, G. Zhong, S. Zhang, M. Li, W. Chen, G. Wang, B. Tu, Y. Wang, X. Chen, B. Ma, S. Li, LML1, Encoding a Conserved Eukaryotic Release Factor 1 Protein, Regulates Cell Death and Pathogen Resistance by Forming a Conserved Complex with SPL33 in Rice. Plant Cell Physiol. 59, 887–902 (2018).

30. V. P. Pisareva, M. A. Skabkin, C. U. T. Hellen, T. V. Pestova, A. V. Pisarev, Dissociation by Pelota, Hbs1 and ABCE1 of mammalian vacant 80S ribosomes and stalled elongation complexes. EMBO J. 30, 1804–1817 (2011).

31. S. Saito, N. Hosoda, S. I. Hoshino, The Hbs1-Dom34 protein complex functions in non-stop mRNA decay in mammalian cells. J. Biol. Chem. 288, 17832–17843 (2013).

32. H. Lange, S. Y. A. Ndecky, C. Gomez-diaz, P. David, N. Butel, J. Zumsteg, L. Kuhn, C. Piermaria, J. Chicher, M. Christie, E. S. Karaaslan, P. L. M. Lang, D. Weigel, H. Vaucheret, P. Hammann, D. Gagliardi, RST1 and RIPR connect the cytosolic RNA exosome to the Ski complex in Arabidopsis. Nat. Commun. 10, 3871 (2019).

33. S. Scholz, J. Pleßmann, B. Enugutti, R. Hüttl, K. Wassmer, K. Schneitz, The AGC protein kinase UNICORN controls planar growth by attenuating PDK1 in Arabidopsis thaliana. PLoS Genet. 15, e1007927 (2019).

34. L. R. Pearce, D. Komander, D. R. Alessi, The nuts and bolts of AGC protein kinases. Nat. Rev. Mol. Cell Biol. 11, 9–22 (2010).

35. H. Zhu, R. G. Kranz, A nitrogen-regulated glutamine amidotransferase (GAT1_2.1) represses shoot branching in Arabidopsis. Plant Physiol. 160, 1770–1780 (2012).

36. Y. Gao, A. A. Badejo, Y. Sawa, T. Ishikawa, Analysis of two L-Galactono-1,4-Lactone-responsive genes with complementary expression during the development of Arabidopsis thaliana. Plant Cell Physiol. 53, 592–601 (2012).

37. N. Tanaka, H. Uno, S. Okuda, S. Gunji, A. Ferjani, T. Aoyama, M. Maeshima, SRPP, a cell wall protein is involved in development and protection of seeds and root hairs in Arabidopsis thaliana. Plant Cell Physiol. 58, 760–769 (2017).

38. A. K. Boron, J. Van Orden, M. N. Markakis, G. Mouille, D. Adriaensen, J. P. Verbelen, H. Höfte, K. Vissenberg, Proline-rich protein-like PRPL1 controls elongation of root hairs in Arabidopsis thaliana. J. Exp. Bot. 65, 5485–5495 (2014).

39. E. Del Campillo, S. Gaddam, D. Mettle-Amuah, J. Heneks, A tale of two tissues: AtGH9C1 is an endo-β-1,4-glucanase involved in root hair and endosperm development in Arabidopsis. PLoS One. 7, e49363 (2012).

40. C. Ringli, N. Baumberger, B. Keller, The Arabidopsis root hair mutants der2-der9 are affected at different stages of root hair development. Plant Cell Physiol. 46, 1046–1053 (2005).

41. R. G. Anthony, R. Henriques, A. Helfer, T. Mészáros, G. Rios, C. Testerink, T. Munnik, M. Deák, C. Koncz, L. Bögre, A protein kinase target of a PDK1 signalling pathway is involved in root hair growth in Arabidopsis. EMBO J. 23, 572–581 (2004).

42. B. P. H. J. Thomma, K. Eggermont, B. Mauch-Mani, R. Vogelsang, B. P. A. Cammue, W. F. Broekaert, Separate jasmonate-dependent and salicylate-dependent defense-response pathways in Arabidopsis are essential for resistance to distinct microbial pathogens. Proc. Natl. Acad. Sci. USA. 95, 15107–15111 (1998).

43. M. Tronchet, C. BalaguÉ, T. Kroj, L. Jouanin, D. Roby, Cinnamyl alcohol dehydrogenases-C and D, key enzymes in lignin biosynthesis, play an essential role in disease resistance in Arabidopsis. Mol. Plant Pathol. 11, 83–92 (2010).

44. C. C. Chen, W. F. Chien, N. C. Lin, K. C. Yeh, Alternative functions of Arabidopsis YELLOW STRIPELIKE3: From metal translocation to pathogen defense. PLoS One. 9, 1–6 (2014).

45. H. U. Stotz, Y. Sawada, Y. Shimada, M. Y. Hirai, E. Sasaki, M. Krischke, P. D. Brown, K. Saito, Y. Kamiya, Role of camalexin, indole glucosinolates, and side chain modification of glucosinolate-derived isothiocyanates in defense of Arabidopsis against Sclerotinia sclerotiorum. Plant J. 67, 81–93 (2011).

46. J. Jung, K. Kumar, H. Y. Lee, Y.-I. Park, H.-T. Cho, S. B. Ryu, Translocation of phospholipase A2α to apoplasts is modulated by developmental stages and bacterial infection in Arabidopsis. Front. Plant Sci. 3 (2012), p. 126.

47. H. H. Breitenbach, M. Wenig, F. Wittek, L. Jorda, A. M. Maldonado-Alconada, H. Sarioglu, T. Colby, C. Knappe, M. Bichlmeier, E. Pabst, D. Mackey, J. E. Parker, A. C. Vlot, Contrasting roles of apoplastic aspartyl protease AED1 and legume lectin-like protein LLP1 in Arabidopsis systemic acquired resistance. Plant Physiol. 165, 791–809 (2014).

48. S. Ferrari, Tandemly duplicated Arabidopsis genes that encode polygalacturonase-inhibiting proteins are regulated coordinately by different signal transduction pathways in response to fungal infection. Plant Cell. 15, 93–106 (2003).

49. S. Ferrari, R. Galletti, D. Vairo, F. Cervone, G. De Lorenzo, Antisense expression of the Arabidopsis thaliana AtPGIP1 gene reduces polygalacturonase-inhibiting protein accumulation and enhances susceptibility to Botrytis cinerea. Mol. Plant-Microbe Interact. 19, 931–936 (2007).

50. C. Weis, U. Hildebrandt, T. Hoffmann, C. Hemetsberger, S. Pfeilmeier, C. König, W. Schwab, R. Eichmann, R. Hückelhoven, CYP83A1 is required for metabolic compatibility of Arabidopsis with the adapted powdery mildew fungus Erysiphe cruciferarum. New Phytol. 202, 1310–1319 (2014).

51. M. Uemura, S. J. Gilmour, M. F. Thomashow, P. L. Steponkus, Effects of COR6.6 and CORl5am polypeptides encoded by COR (Cold-Regulated) genes of Arabidopsis thaliana on the freeze-induced fusion and leakage of liposomes. Plant Physiol. 111, 313–327 (2002).

52. D. P. Horvath, B. K. Mclarney, M. F. Thomashow, Regulation of Arabidopsis thaliana L. (Heyn) cor78 in response to low temperature. Plant Physiol. 103, 1047–1053 (1993).

53. B. C. Dyson, M. A. E. Miller, R. Feil, N. Rattray, C. G. Bowsher, R. Goodacre, J. E. Lunn, G. N. Johnson, FUM2, a cytosolic fumarase, is essential for acclimation to low temperature in Arabidopsis thaliana. Plant Physiol. 172, 118–127 (2016).

54. L. Guo, H. Yang, X. Zhang, S. Yang, Lipid transfer protein 3 as a target of MYB96 mediates freezing and drought stress in Arabidopsis. J. Exp. Bot. 64, 1755–1767 (2013).

55. X. B. Zhang, B. H. Feng, H. M. Wang, X. Xu, Y. F. Shi, Y. He, Z. Chen, A. P. Sathe, L. Shi, J. L. Wu, A substitution mutation in OsPELOTA confers bacterial blight resistance by activating the salicylic acid pathway. J. Integr. Plant Biol. 60, 160–172 (2018).

56. T. A. Masters, V. Calleja, D. A. Armoogum, R. J. Marsh, C. J. Applebee, M. Laguerre, A. J. Bain, B. Larijani, Regulation of 3-phosphoinositide-dependent protein kinase 1 activity by homodimerization in live cells. Sci. Signal. 3 (2010), doi:10.1126/scisignal.2000738.

57. I. M. Adham, M. A. Sallam, G. Steding, M. Korabiowska, U. Brinck, S. Hoyer-fender, C. Oh, W. Engel, Disruption of the Pelota Gene Causes Early Embryonic Lethality and Defects in Cell Cycle Progression. Mol. Cell. Biol. 23, 1470–1476 (2003).

58. M. M. Mahfouz, Arabidopsis TARGET OF RAPAMYCIN Interacts with RAPTOR, Which Regulates the Activity of S6 Kinase in Response to Osmotic Stress Signals. Plant Cell. 18 (2006), pp. 477–490.

59. X. Chen, S. M. Goodwin, X. Liu, X. Chen, R. A. Bressan, M. A. Jenks, Mutation of the RESURRECTION1 locus of Arabidopsis reveals an association of cuticular wax with embryo development. Plant Physiol. 139, 909–919 (2005).

60. T. Li, A. Natran, Y. Chen, J. Vercruysse, K. Wang, N. Gonzalez, M. Dubois, D. Inzé, A genetics screen highlights emerging roles for CPL3, RST1 and URT1 in RNA metabolism and silencing. Nat. Plants. 5, 539–550 (2019).

61. Q. Zhao, J. Shen, C. Gao, Y. Cui, Y. Wang, J. Cui, L. Cheng, W. Cao, RST1 is a FREE1 suppressor that negatively regulates vacuolar rrafficking in Arabidopsis. Plant Cell (2019), doi:10.1105/tpc.19.00003.

62. B. M. Waters, H.-H. Chu, R. J. DiDonato, L. A. Roberts, R. B. Eisley, B. Lahner, D. E. Salt, E. L. Walker, Mutations in Arabidopsis Yellow Stripe-Like1 and Yellow Stripe-Like3 reveal their roles in metal ion homeostasis and loading of metal ions in seeds. Plant Physiol. 141, 1446–1458 (2006).

63. T. Shimada, K. Yamada, M. Kataoka, S. Nakaune, Y. Koumoto, M. Kuroyanagi, S. Tabata, T. Kato, K. Shinozaki, M. Seki, M. Kobayashi, M. Kondo, M. Nishimura, I. Hara-Nishimura, Vacuolar processing enzymes are essential for proper processing of seed storage proteins in Arabidopsis thaliana. J. Biol. Chem. 278, 32292–32299 (2003).

64. Q. Li, B.-C. Wang, Y. Xu, Y.-X. Zhu, Systematic studies of 12S seed storage protein accumulation and degradation patterns during Arabidopsis seed maturation and early seedling germination stages. BMB Rep. 40, 373–381 (2011).

65. X. Tang, M. H. Lim, J. Pelletier, M. Tang, V. Nguyen, W. A. Keller, E. W. T. Tsang, A. Wang, S. J. Rothstein, J. J. Harada, Y. Cui, Synergistic repression of the embryonic programme by SET DOMAIN GROUP 8 and EMBRYONIC FLOWER 2 in Arabidopsis seedlings. J. Exp. Bot. 63, 1391–1404 (2012).

66. M. K. Choy, J. A. Sullivan, J. C. Theobald, W. J. Davies, J. C. Gray, An Arabidopsis mutant able to green after extended dark periods shows decreased transcripts of seed protein genes and altered sensitivity to abscisic acid. J. Exp. Bot. 59, 3869–3884 (2008).

67. T. Umezawa, Y. Fujita, T. Furihata, K. Maruyama, K. Yamaguchi-Shinozaki, K. Shinozaki, R. Yoshida, Abscisic acid-dependent multisite phosphorylation regulates the activity of a transcription activator AREB1. Proc. Natl. Acad. Sci. 103, 1988–1993 (2006).

68. L. Li, T. Shimada, H. Takahashi, H. Ueda, Y. Fukao, M. Kondo, M. Nishimura, I. Hara-Nishimura, MAIGO2 is involved in exit of seed storage proteins from the endoplasmic reticulum in Arabidopsis thaliana. Plant Cell. 18, 3535–3547 (2006).

69. K. Müntz, Deposition of storage proteins. Plant Mol. Biol. 38, 77–99 (1998).

70. J. M. Alonso, T. Curran, R. Hawkes, P. Soriano, J. A. Cooper, J. W. Lichtman, B. Bernier, A. M. Goffinet, M. Derer, A. Goffinet, M. J. Galazo, C. Cavada, J. A. Conchello, L. T. Landmesser, R. D. Fields, B. W. Festoff, P. G. Nelson, I. V Smirnova, B. A. Citron, Genome-wide insertional mutagenesis of Arabidopsis thaliana. Science. 301, 653–657 (2003).

71. S. J. Clough, A. F. Bent, Floral dip: A simplified method for Agrobacterium-mediated transformation of Arabidopsis thaliana. Plant J. 16, 735–743 (1998).

72. W. Liu, Z. H. Xu, D. Luo, H. W. Xue, Roles of OsCKI1, a rice casein kinase I, in root development and plant hormone sensitivity. Plant J. 36, 189–202 (2003).

73. S.-T. Tan, H.-W. Xue, Casein Kinase 1 Regulates Ethylene Synthesis by Phosphorylating and Promoting the Turnover of ACS5. Cell Rep. 9 (2014), doi:10.1016/j.celrep.2014.10.047.

74. S. Nakamura, S. Mano, Y. Tanaka, M. Ohnishi, C. Nakamori, M. Araki, T. Niwa, M. Nishimura, H. Kaminaka, T. Nakagawa, Y. Sato, S. Ishiguro, Gateway binary vectors with the bialaphos resistance gene, bar, as a selection marker for plant transformation. Biosci. Biotechnol. Biochem. 74, 1315–1319 (2010).

75. B. K. Nelson, X. Cai, A. Nebenführ, A multicolored set of in vivo organelle markers for co-localization studies in Arabidopsis and other plants. Plant J. 51, 1126–1136 (2007).

76. D. Lin, S. Nagawa, J. Chen, L. Cao, X. Chen, T. Xu, H. Li, P. Dhonukshe, C. Yamamuro, J. Friml, B. Scheres, Y. Fu, Z. Yang, A ROP GTPase-dependent auxin signaling pathway regulates the subcellular distribution of PIN2 in Arabidopsis roots. Curr. Biol. 22, 1319–1325 (2012).

77. B. J. Yang, X. X. Han, L. L. Yin, M. Q. Xing, Z. H. Xu, H. W. Xue, Arabidopsis PROTEASOME REGULATOR1 is required for auxin-mediated suppression of proteasome activity and regulates auxin signalling. Nat. Commun. 7, 11388 (2016).

78. J. Zhang, T. Nodzynski, A. Pencik, J. Rolcik, J. Friml, PIN phosphorylation is sufficient to mediate PIN polarity and direct auxin transport. Proc. Natl. Acad. Sci. 107, 918–922 (2010).

79. A. Thompson, J. Schäfer, K. Kuhn, S. Kienle, J. Schwarz, G. Schmidt, T. Neumann, R. Johnstone, A. K. A. Mohammed, C. Hamon, Tandem mass tags: a novel quantification strategy for comparative analysis of complex protein mixtures by MS/MS. Anal. Chem. 75 (2003), pp. 1895–904.

80. J. Schindelin, I. Arganda-Carreras, E. Frise, V. Kaynig, M. Longair, T. Pietzsch, S. Preibisch, C. Rueden, S. Saalfeld, B. Schmid, J. Y. Tinevez, D. J. White, V. Hartenstein, K. Eliceiri, P. Tomancak, A. Cardona, Fiji: An open-source platform for biological-image analysis. Nat. Methods. 9, 676–682 (2012).

